# High-density CRISPRi screens reveal adaptive transcriptional gradients in cyanobacteria

**DOI:** 10.1101/2024.05.20.595006

**Authors:** Andrew Hren, Nicole Lollini, Dana L. Carper, Paul Abraham, Jeffrey C. Cameron, Jerome M. Fox, Carrie Eckert

**Affiliations:** Department of Chemical and Biological Engineering, University of Colorado, Boulder, CO, 80304, USA; Biosciences Division, Oak Ridge National Laboratory, Oak Ridge, TN, 37831, USA; Department of Biochemistry, University of Colorado, Boulder, CO, 80309, USA; Renewable and Sustainable Energy Institute, University of Colorado, Boulder, CO, 80309, USA; National Renewable Energy Laboratory, Golden, CO, 80401, USA

**Keywords:** genome-wide screens, environmental adaptation, spectral tuning, photosynthesis, energy transduction

## Abstract

Cyanobacteria are the oldest form of photosynthetic life on Earth and contribute to primary production in nearly every habitat, from permafrost to hot springs. Despite longstanding interest in the biochemical basis of environmental adaptation in these microbes, it remains poorly understood and challenging to re-wire. This study uses a high-density, genome-wide CRISPR interference (CRISPRi) screen to examine the influence of gene-specific transcriptional variation on the growth of *Synechococcus* sp. PCC 7002 under environmental extrema. Surprisingly, many partial knockdowns enhanced fitness under cold monochromatic conditions. Notably, transcriptional repression of a gene for a core subunit of the NDH-1 complex, which is important for photosynthesis and carbon uptake, improved growth rates under both red and blue light but at distinct, color-specific optima. In general, most genes with fitness-improving knockdowns were distinct to each light color, evidencing unique stress responses and alleviation mechanisms. Multi-target transcriptional repression produced nonadditive effects. Findings reveal diverse mechanisms of environmental adaptation in cyanobacteria and provide a new approach for using gradients in sgRNA activity to pinpoint biochemically influential transcriptional changes in cells.

**SIGNIFICANCE STATEMENT:** Cyanobacteria are the most abundant photosynthetic organisms on Earth, where they endure a striking variety of environmental fluctuations. This study examines the biochemical basis of environmental adaptation in *Synechococcus* sp. PCC 7002, an important model strain, by modulating the expression of every gene in its genome. Results show that partial, but not complete, reduction in the expression of a subset of influential genes can improve growth under cold monochromatic conditions. Optimal expression levels differ between red and blue light and shift with multi-gene adjustments. Findings show how minor transcriptional adjustments can yield major improvements in growth under environmental extrema and provide a powerful systems-level approach for studying—and fine-tuning—the adaptive capacity of microbes.

## INTRODUCTION

Cyanobacteria, like all cellular systems, rely on a limited set of dynamically regulated biochemical pathways to grow in environments that change, often irregularly, over time and space (1). Shifts in intracellular protein concentration caused by stimuli-responsive transcriptional circuits play a critical role in enabling their adaptation to diverse light, temperature, and nutrient regimes (2). In one prominent example, the circadian clock creates regular shifts in protein phosphorylation to regulate rhythmic gene expression over 24-hour periods, adjusting protein concentration in anticipation of periodic environmental shifts caused by the Earth’s rotation (3). Cyanobacteria can also respond to the color of light, or spectral quality. Mechanisms of chromatic acclimation vary by species, but all involve light-responsive transcriptional circuits that change the stoichiometry of core photosynthetic components to improve their spectral compatibility with incident wavelengths (1). Responses to temperature, in turn, rely on changes in the expression of fatty acid desaturases (4) that adjust membrane fluidity. In general, intermediate transcriptional adjustments are a hallmark of adaptation in cyanobacteria (5, 6), and new methods for surveying their influence on growth under diverse environmental conditions could improve our understanding of—and ability to engineer—their adaptive responses.

Contemporary approaches to survey gene function tend to focus on conditional fitness. For cyanobacteria, prominent examples include gene inactivation via transposon mutagenesis (TnSeq) (7) and transcriptional repression via CRISPR interference (CRISPRi) (8, 9). Both methods can leverage large libraries of mutants with targeted, sequence-trackable perturbations (10). When paired with biological selection and massively parallel sequencing, in turn, they can uncover genes that affect growth under distinct environmental conditions (e.g., light cycles or media compositions) (9). To date, most studies of fitness have focused on generating gene-specific classifiers, for example, essential, non-essential, or advantageous (7, 11, 12). While valuable, these classifiers overlook the transcriptional dependence of fitness effects—a dependency consistent with the core biochemical mechanisms that underly environmental adaptation in cyanobacteria.

CRISPRi can generate intermediate levels of transcriptional repression and, thus, provides a powerful—if often overlooked—tool for examining the influence of systematically titrated biochemical adjustments on biological adaptation. The most widely used CRISPRi system uses a mutationally inactivated Cas9 endonuclease (dCas9) to bind target genes and interfere with transcription; the sequence of the single-guide RNA (sgRNA), which complexes with dCas9, determines the location of binding and the strength of inhibition (13). In this study, we used high-density sgRNA libraries (∼10 guides per gene) to carry out a genome-wide analysis of single-gene transcriptional gradients in *Synechococcus* sp. PCC 7002 (PCC 7002), a fast-growing and industrially relevant model cyanobacterium (14–16). By examining changes in the composition of these libraries (i.e., sgRNA enrichment) after growth under environmental extrema, we identified genes for which intermediate levels of transcriptional repression improved growth. Strikingly, at low temperatures, repression of a gene for a core subunit of NDH-1, a broadly influential complex, improved growth under both red and blue light, but at distinct transcriptional optima. In general, the effects of intermediate levels of transcriptional repression shifted with illumination intensity and exhibited super- and subadditivity in dual-guide systems, effects that are lost—and nearly impossible to predict—with common gene-specific classifiers (i.e., essential or non-essential).

## RESULTS

### Genome-wide CRISPRi screens reveal condition-specific enrichment of sgRNAs

The genome of PCC 7002 contains 3237 genes, of which ∼35% are unannotated (17). To support genome-wide CRISPRi screens, we prepared cells with genomically integrated copies of dCas9 and sgRNAs under the control of inducible promoters (Fig. 1A). For each gene, we designed multiple sgRNAs (∼10 per gene) by using a custom software package to maximize the uniqueness of the seed sequence (i.e., the first 7-12 nucleotides at the 3′ end of the sgRNA spacer neighboring the protospacer adjacent motif, or PAM) and minimize the distance between this sequence and the start codon on the coding strand—properties that enhance transcriptional repression (SI Appendix, Figs. S1A-S1C; (9, 18, 19)). The final gene-specific sgRNAs met these design criteria to different degrees, exhibited natural variability in their PAM-proximal sequence composition, or had different predicted secondary structures—established sources of variability in transcriptional repression (SI Appendix, Figs. S1D and S2; (20)). In the final library, 98.8% of genes had 10 sgRNAs with variable strengths (SI Appendix, Fig. S3).

**Figure 1.**
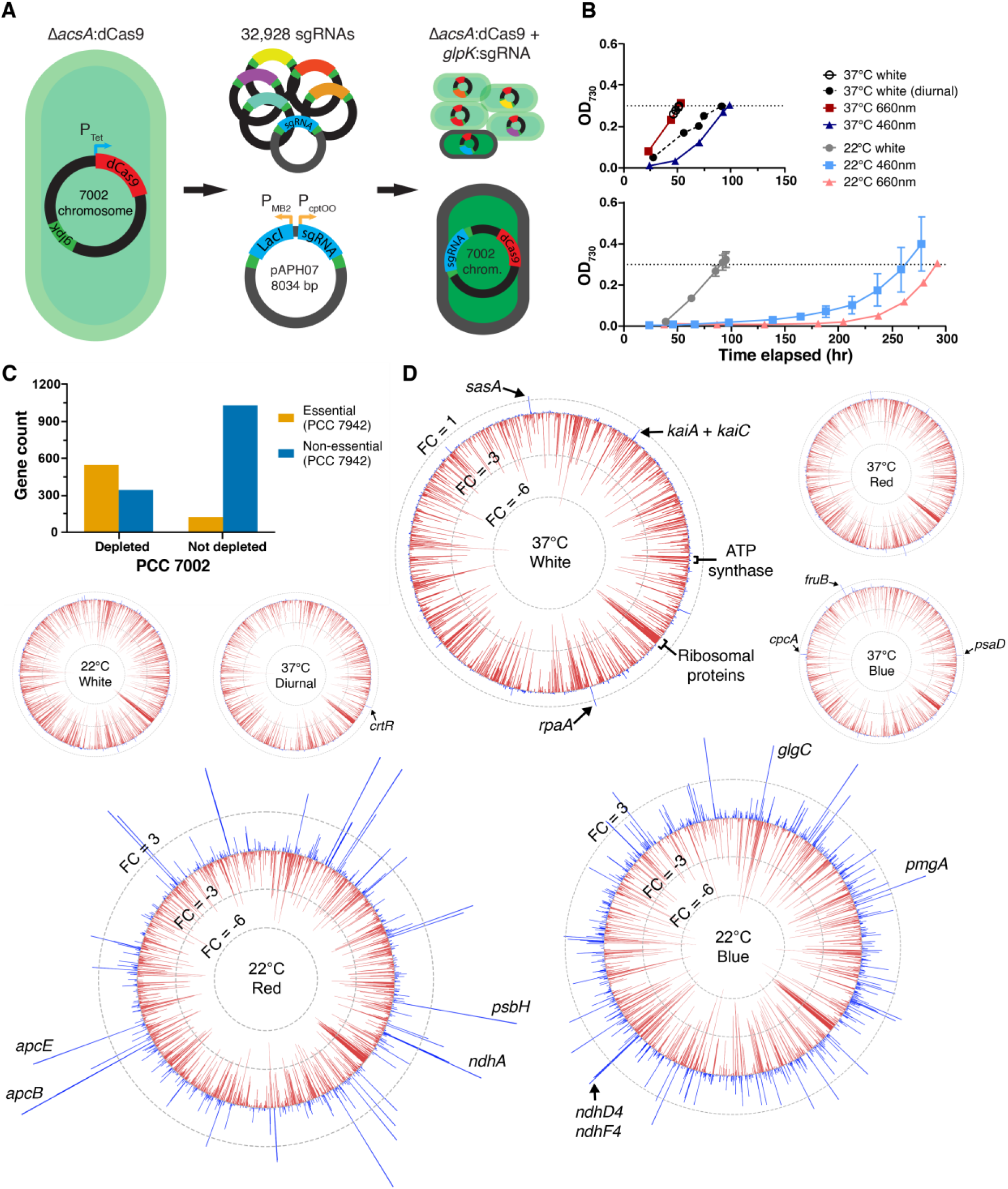
Genome-wide CRISPRi screens reveal condition-specific enrichment of sgRNAs in PCC 7002. (A) For each screen, we transformed a strain of PCC 7002 with dCas9 under the control of an anhydrotetracycline-inducible promoter (*ΔacsA*:dCas9) with a plasmid-borne sgRNA library, which placed each sgRNA downstream of *glpK* under the control of an IPTG-inducible promoter. (B) We grew transformed cells under seven growth conditions that differed in temperature (i.e., 22 or 37°C) or light regime (i.e., color or exposure duration), harvested cells at an OD_730_ of at least 0.3 (∼9-10 doublings). (C) We used next-generation sequencing (NGS) to compare the abundance of sgRNAs before and after growth. The plot enumerates genes with or without significant sgRNA depletion in at least one condition (“depleted” or “not depleted”) and compares them with essentiality designations from a prior Tn-Seq analysis of *S. elongatus* (7). (D) Genome maps showing the mean sgRNA enrichment for growth under each condition; bar positioning matches the chromosomal location of each gene. Cold, narrow-spectrum conditions reveal numerous growth-enhancing knockdowns (blue bars). Data in panels (B) and (D) represent the mean and SD of n=2 biological replicates. Table S1 provides all sgRNA enrichment values.

Many environmental stresses exhibit nonlinear effects when applied in combination (21). To explore this nonlinearity, we grew our cell libraries under white, red, and blue light at both 22°C and 37°C; diurnal white light (37°C) served as a “natural” extremum (22). At 37°C, pooled cells grew more rapidly under white and red light than under blue or diurnal light (Fig. 1B). At 22°C, growth slowed for all light conditions, but most prominently for red and blue, where cells grew approximately three-fold slower than in sustained white light. The profound slowdown in cold monochromatic light suggests that PCC 7002 is ill-equipped to adapt to these conditions.

We began our analysis with a coarse classification of genes based on sgRNA depletion. In brief, we used next-generation sequencing (NGS) to measure changes in sgRNA distributions caused by growth under each condition, a hierarchical mixture model to identify genes with significant shifts in sgRNA frequency (12), and compared our enrichrmnet results to prior work. To begin, we examined the essentiality designations from a Tn-Seq analysis of *Synechococcus elongatus* PCC 7942 (PCC 7942) (i.e., protein homologs; Fig. 1C) (7). For genes with sgRNA depletion in at least one condition, an indication of a fitness defect,over half were deemed essential in the study of PCC 7942, while many (∼39%) were non-essential. For genes with no sgRNA depletion, including those with enrichment, a small fraction (∼11%) were classified as essential in the PCC 7942 study, but the majority were non-essential. We observed similar inconsistencies with a more recent genome-wide CRISPRi screen of *Synechocystis* sp. PCC 6803 (9) (PCC 6803; SI Appendix, Fig. S5). Inconsistencies between studies may result from the conditional essentiality of genes under different growth conditions, incomplete knockdown by CRISPRi, or differences in the functional redundancy of genes between species (i.e., the number of paralogs).

To investigate the conditional essentiality of genes, we mapped the mean enrichment of gene-specific sgRNAs onto the genome (Fig. 1D). In general, influential genes were broadly distributed and most showed sgRNA depletion. Prominent examples include genes for ribosomal proteins and ATP production, which are broadly important for growth, as reported in the recent CRISPRi study of PCC 6803 (9). Intriguingly, a subset of genes showed sgRNA enrichment. For growth under white light at 37°C, this effect was small; notably, sgRNAs for clock proteins (e.g., KaiA, KaiC, SasA, and RpaA) improved fitness, a result that corresponds to observed increases in growth rate from arrhythmic *kaiC* and *sasA* mutants in PCC 7942 (23, 24). For growth under red or blue light at 22°C—hereafter, 22R and 22B—sgRNA enrichment reached extreme levels, an indication that targeted knockdowns can improve growth. In addition to its genome, PCC 7002 has six endogenous plasmids, which showed similar but less dramatic trends in sgRNA enrichment—an effect that may reflect reduced sensitivity to CRISPRi repression at higher copy numbers or, perhaps, a lower abundance of critical proteins, relative to the genome (SI Appendix, Fig. S6) (25).

### Cold conditions elicit divergent transcriptional sensitivities to blue and red light

We began our analysis of fitness-improving knockdowns by examining a subset of well-studied genes associated with photoperception, photosynthesis, and cold shock (Fig. 2A). In general, 22R and 22B conditions brought about the most distinct enrichment patterns. Genes associated with photosystem II (PSII) and the phycobilisome (*apcE* and *apcB*), for example, exhibited pronounced sgRNA enrichment under red light, but not blue, while those related to photosystem I (PSI) and adaptation to cold or high intensity light became enriched only in blue light. Genes associated with circadian rhythm and NDH-1, an 11-protein complex with co-existing variants involved in photosynthesis, respiration, and CO_2_ uptake (26), showed enrichment under both conditions, but the extent of enrichment varied with light color.

**Figure 2.**
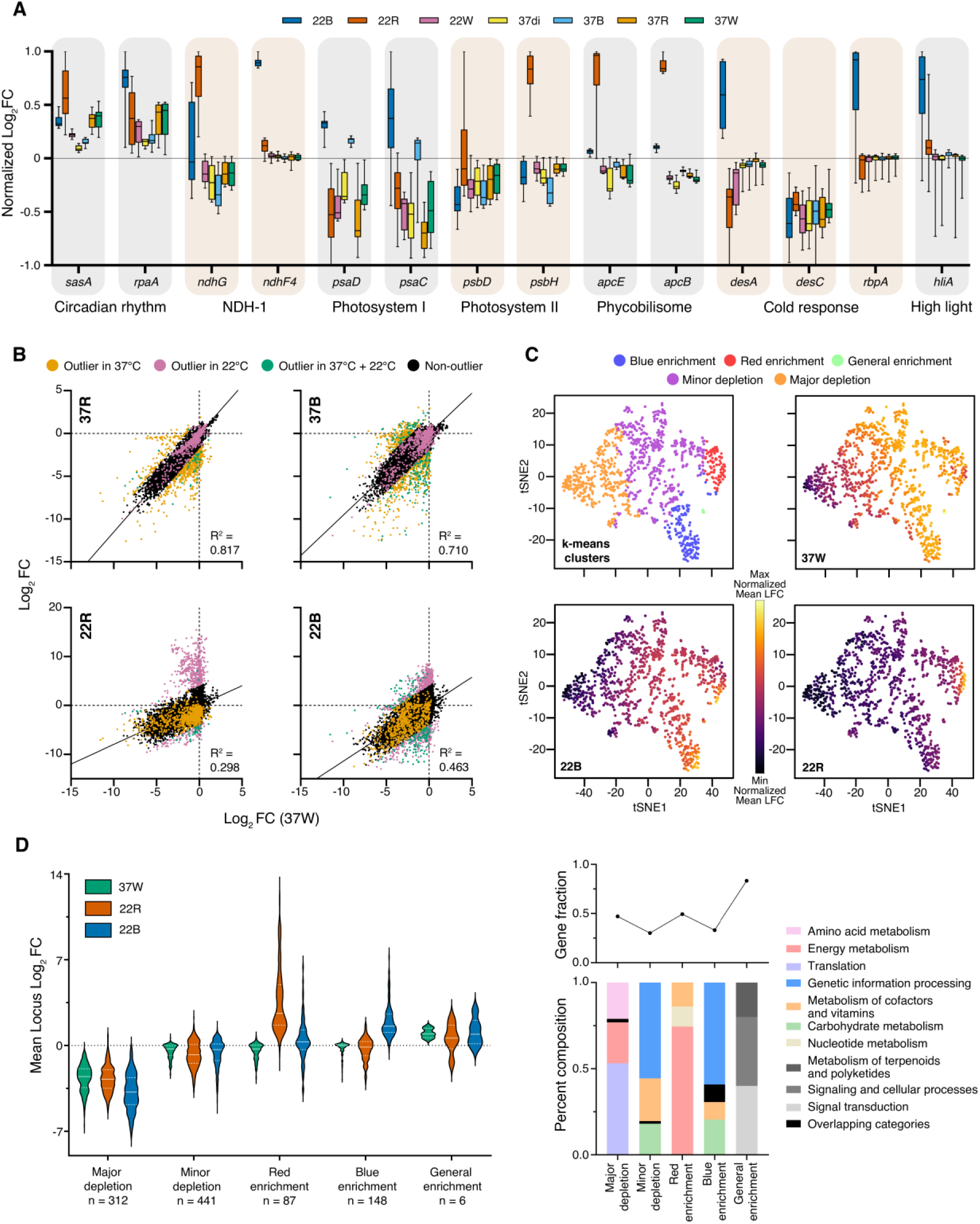
Cold conditions elicit divergent transcriptional sensitivities to blue and red light. (A) The sgRNAs for genes associated with light harvesting, photosynthesis, and cold shock exhibited condition-specific enrichment patterns that diverged most significantly between 22B and 22R. Box plots show the median, upper and lower quartile, and minimum and maximum values of normalized sgRNA enrichment. (B) We analyzed the correlation in sgRNA enrichment between standard growth (37W) and monochromatic conditions. Outlier populations (i.e., sgRNAs with a standardized residual greater than 3) differ between temperatures within each light color. (C) We clustered genes into five groups using enrichment data from environmental extrema (37W, 22R, and 22B) as input parameters in k-means clustering. These clusters, which are plotted here with t-SNE, show condition-specific enrichment patterns. (D) We classified clusters by trends in mean sgRNA enrichment. Violin plots show the median, upper and lower quartile, and minimum and maximum values of mean locus Log_2_FC. (E) The three major gene ontology (GO) categories for each cluster, which account for 30-80% of their constituent genes, reveal functional differences between them. Data depict (A) normalized mean enrichment, (B) DESeq2-calculated sgRNA enrichment, and (D) mean locus enrichment for n=2 biological replicates. Table S1 provides all sgRNA enrichment values.

When examining the influence of monochromatic light, one might naturally ask if a low temperature (22°C) merely enhances stress responses that appear at high temperatures (37°C). A comparison of enrichment between 22°C and 37°C, however, suggests that it does not. Correlations in sgRNA enrichment between (i) white light at 37°C and (ii) red or blue light at 22°C or 37°C show temperature-specific differences in outlier populations (Fig. 2B). This discrepancy is particularly prominent under red light. The outliers indicate that the fitness effects of knockdowns under cold monochromatic conditions are complex and do not reflect a sum of the fitness effects that appear when light or temperature are changed in isolation.

We used a clustering analysis to place genes into five groups with shared fitness effects under 22R, 22B, and 37°C white (37W). When plotted with t-distributed stochastic neighbor embedding (t-SNE), a common method for dimensionality reduction (27), clusters showed condition-specific patterns (Fig. 2C). We classified these clusters by trends in mean sgRNA enrichment: major depletion, minor depletion, red enrichment, blue enrichment, and general enrichment (Fig. 2D). The “depletion” and “general enrichment” groups include genes with condition-independent fitness effects; the red and blue groups, genes with sgRNAs enriched in 22R or 22B.

We extracted the three most common gene ontology (GO) categories for each cluster (Fig. 2E). Genes associated with translation showed sgRNA depletion across conditions (“major depletion”), a finding consistent with previously reported fitness defects caused by the repression of ribosomal genes (28). Meanwhile, genes for signaling proteins (e.g., the Kai oscillator) were broadly enriched, an indication that many sensory proteins are no longer necessary when light is constant, regardless of spectral quality. For both “minor depletion” and “blue enrichment”, in turn, genes for genetic information processing (e.g., DNA transcription, replication, and repair) were most common; accordingly, some mild, yet generally unfavorable biochemical changes may become favorable under 22B. Finally, for 22R, energy metabolism (e.g., *apc*, *ndh*, and *psb*) was the largest subgroup. Overall, the lack of major GO overlap between clusters suggests that stress responses and alleviation mechanisms are distinct to each environmental extrema.

### Enrichment gradients reveal intermediate, fitness-improving knockdowns

Differences in sgRNA enrichment between growth conditions illustrate how a gene can appear neutral in one environment but highly influential in others (Fig. 2A). We observed many examples of this context dependence (SI Appendix, Fig. S7). For example, several genes with gradients in sgRNA depletion under 37W exhibited both depletion and enrichment under 22B and 22R (Fig. 3A). The *ndhG* gene provides a striking example of condition-specific optima. This gene encodes a core structural component of NDH-1. The sgRNA enrichment levels for *ndhG* showed a linear correlation between 37W, where all sgRNAs were depleted (Log_2_FC = -2 to 0), and 22B, where several sgRNAs were highly enriched (Log_2_FC = -3 to 5.5; Fig 3B). Under 22R, the same set of sgRNAs conferred a broad fitness improvement with an intermediate optimum. We speculated that gradients in sgRNA depletion under 37W might correspond to gradients in transcriptional repression. To test this theory, we expressed three *ndhG* guides with different levels of depletion and measured mRNA levels (Fig. 3C). Indeed, transcript levels correlated with sgRNA depletion at 37W, suggesting that this change provides a reasonable metric for guide strength; gradients in sgRNA enrichment under environmental extrema thus reveal adaptive transcriptional gradients. Differential guide enrichment under 22R and 22B, in turn, indicates that medium and low transcriptional repression of the *ndhG* gene, respectively, improve adaptation to these conditions.

**Figure 3.**
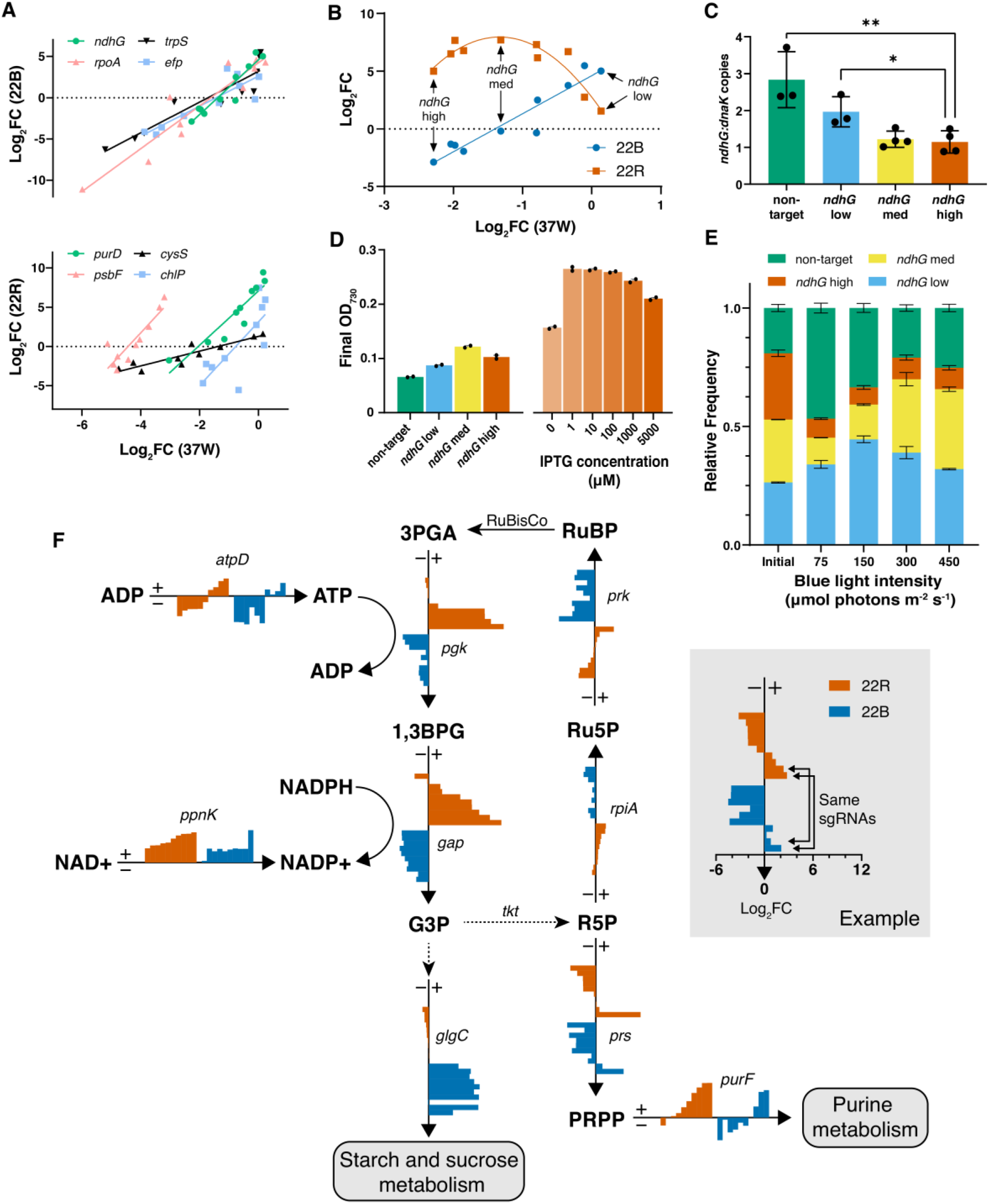
Intermediate transcriptional repression can improve fitness under cold, narrow spectrum conditions. (A) In 37W, the sgRNAs for a subset of genes exhibited a broad range of depletion, an indication of differences in guide strength; in 22B and 22R, the same sgRNAs showed correlating levels of depletion and enrichment. (B) Correlations associated with *ndhG* changed between light conditions. (C-E) Analysis of *ndhG*. (C) For three guides, the extent of sgRNA depletion under 37W correlates with transcriptional repression (i.e., mRNA levels) in PCC 7002. (D) We examined the influence of *ndhG* on growth under 22B by using (left) sgRNAs with different strengths (i.e., guides from C) or (right) a single sgRNA under different levels of induction. In both experiments, intermediate transcriptional repression of *ndhG* improved fitness (i.e., increased cell density). (E) In a focused library of sgRNAs with different strengths, frequencies at 22°C changed with blue light intensity relative to the initial distribution. (F) Genes within central metabolic pathways display disparate fitness profiles and gradients between growth conditions. Here, we sorted all sgRNAs by the Log_2_FC values under 22R. Data in (A), (B), and (F) depict the Log_2_FC of individual sgRNAs. Data in C-E depict (C), the mean and standard deviation for n ≥3 technical replicates of n = 2 biological replicates and D-E the mean and standard error for n = 2 biological replicates. Statistics were performed using two-tailed *t*-tests. (\**P*<0.05 and \*\**P*<0.01)

We examined *ndhG* as a representative “gradient gene” by using orthogonal experiments to confirm its influence on fitness under 22B. First, we measured the growth of PCC 7002 strains containing low-, medium-, or high-strength sgRNAs; indeed, the intermediate-strength sgRNA improved growth (Fig. 3D). Second, we used IPTG to tune the expression of the strong sgRNA; here, a low level of induction enhanced fitness under 22B. The discrepancy in apparent optima between experiments (i.e., low repression vs. high) could reflect differences in the mRNA levels afforded by each transcriptional control strategy or, perhaps more simply, differences in light intensity. Indeed, a comparison of light intensities showed a shift in optimal guide strength as intensity increased (Fig. 3E). In general, the influence of light intensity highlights the value of using high-density sgRNA libraries to identify fitness-improving knockdowns, which might become apparent within narrow, highly sensitive windows of transcriptional repression.

Genes that showed gradients in sgRNA enrichment tended to be essential, based on a prior analysis of PCC 7942 (7). The Calvin-Benson-Bassham (CBB) cycle required for CO_2_ fixation provides a representative example (Fig. 3F; see Table S6 for gene products). Under both 22R and 22B, *purF* and *atpD* shared similar sgRNA gradients, which suggest that a slight reduction in purine metabolism and ATP production are broadly beneficial. The effect of *purF* is consistent with prior work on purine biosynthesis, which is disfavored under energy stress (29, 30). Any knockdown of *ppnK*, which should increase the NAD+/NADP+ ratio (31), also improved fitness under both conditions. In 22B, knockdown of *glgC* is uniquely advantageous. This gene is commonly repressed in metabolic engineering efforts to increase carbon flux toward target metabolites (32, 33). Our analysis illustrates how mapping gradient effects onto essential pathways can reveal subtle changes in transcription that can improve fitness under extreme conditions.

### Cold monochromatic conditions create an imbalance between photosystems I and II

Extremely cold temperatures enable resolution of the fluorescence emission of light-harvesting systems. We used low-temperature (77 K) fluorescence spectroscopy to examine PSI, PSII, and their coupling phycobilisomes (PBS), light-harvesting antenna that interact with both photosystems to rebalance excitation energy. First, we excited chlorophyll *a* (λ_ex_ = 440 nm) to measure relative amounts of PSI (λ_em_ = 720 nm) and PSII (λ_em_ = 685/695 nm; Fig. 4A) (34, 35). The PSI:PSII ratio decreased substantially under 22B relative to 22W but remained unchanged in 22R (SI Appendix, Fig. S8). Next, we excited phycocyanin (λ_ex_ = 580 nm) to measure changes in the distribution of PBS (Fig. 4B) (36). Blue light increased the proportion of both free PBS and PSII-PBS complex, relative to their levels under red and white light. These shifts are similar to those observed in PCC 6803, which grows slowly and exhibits low oxygen production in blue light (37). This poor photosynthetic performance has been attributed to the overexcitation of PSI, which has three-fold more chlorophyll *a*—and more blue-light absorbing β-carotenes—than PSII (38). In both PCC 6803 and PCC 7002, cells respond to this photosystem imbalance by coupling PBSs to PSII (state 1), a shift that can increase the absorption cross-section of PSII and enhance linear electron transport (LET); PBSs, however, cannot absorb blue light effectively (37), so the compensatory effect is limited. One might expect a similar imbalance under red light, where PSI still benefits from a high chlorophyll content, but PBSs absorb red light effectively.

**Figure 4.**
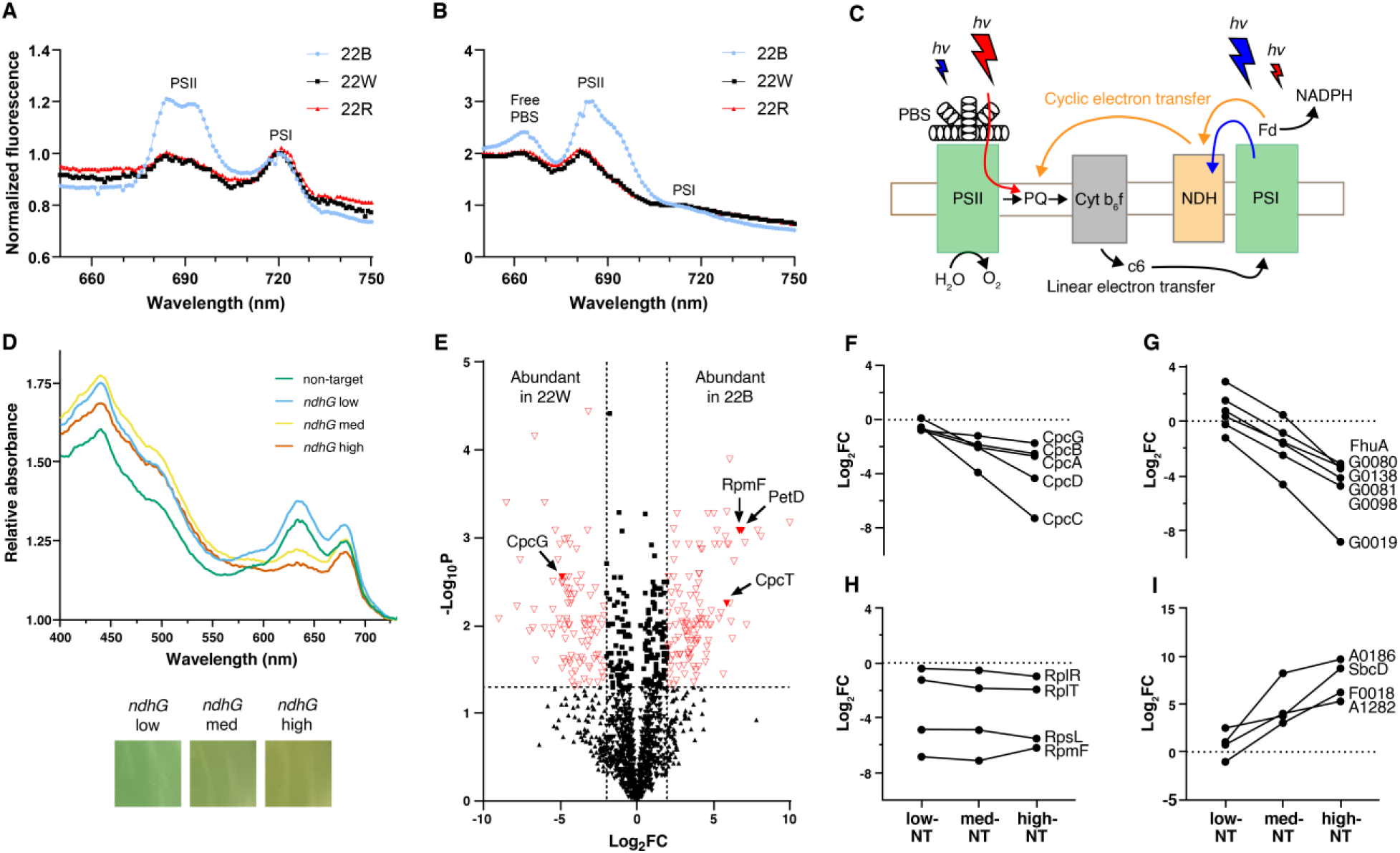
Cold monochromatic conditions induce stress responses that can be mitigated or enhanced by different levels of *ndhG* repression. (A-B) Low-temperature (77 K) fluorescence emission spectra of PCC 7002 cultures exposed to blue (460 nm), red (660 nm), or white light at 22℃ (22B, 22R, and 22W, respectively). (A) Excitation of chlorophyll *a* at 440 nm yields fluorescence emission peaks for PSI (λ_em_ = 720 nm) and PSII (λ_em_ = 685/695 nm) (34, 35); spectra are normalized to 720 nm. The 22B conditions decrease the PSI:PSII ratio, relative to white and red light. (B) Excitation of phycocyanin at 580 nm yields fluorescence emission peaks for free PBS (660 nm), PBS-PSII (685 nm), and PBS-PSI (712 nm). Spectra are normalized to PBS-PSI (712 nm). The 22B condition increases the proportion of free PBS and PBS-PSII complex. (C) A schematic of photosystem imbalance. At 22℃, blue light overexcites PSI (37) and enhances PQ oxidation (49), which is mitigated by NDH-1 through CET. Red light overexcites PSII and enhances PQ reduction through linear electron transport (LET). (D) Absorbance spectra of cells grown in 22B. Minor repression of *ndhG* enhances chlorophyll a and phycocyanin abundance; higher levels of repression decrease this effect. (E) Many proteins exhibit significant differences in abundance (red, p < 0.05, |Log_2_FC| > 2) between 22B and 22W in the non-targeting strain. (F-I), Analysis of changes in proteins under 22B. Increasing levels of *ndhG* repression decrease the relative abundance of both (F) core phycobilisome proteins and (G) iron-handling proteins, (H) leave ribosomal proteins at similar levels of depletion, and (I) increase the relative abundance of DNA repair and uncharacterized proteins. Data in E-I represent the normalized, mean-centered Log_2_FC of 4 biological replicates. Full protein names appear in Table S2.

### Repression of *ndhG* enhances general stress responses under 22B

In cyclic electron transport (CET), NDH-1 accepts electrons from the acceptor side of PSI (39) and transfers them to the plastiquinone (PQ) pool, enabling the production of ATP without NADPH (40) to supplement the insufficient ATP produced via LET for cellular metabolism (41). By attenuating this essential process, repression of *ndhG* has the potential to be toxic. Indeed, gradients in sgRNA enrichment under 22B conditions suggest that high repression is deleterious. To investigate this effect further, we monitored the influence of *ndhG* repression on the production of light-harvesting pigments under 22B. Low-strength sgRNA increased absorption peaks for both chlorophyll a (440 and 680 nm) and phycocyanin (620 nm), while the medium- and high-strength sgRNAs decreased them (Fig. 4D). These changes suggest that minor *ndhG* repression enhances the native cellular response to 22B by increasing the proportion of PSII and PSII-PBS complexes. High repression mutes this response, perhaps by enhancing cellular stress.

We broadened our analysis of *ndhG* repression by using quantitative proteomics. Interestingly, we could not reliably detect NdhG—a potential reflection of its low abundance (other NDH-1 subunits gave low signals) or poor compatibility with the method—but many proteins showed major shifts between 22W and 22B (e.g., PBS proteins; Fig. 4E). We focused our analysis on proteins that showed different trends with *ndhG* repression. Under 22B, the abundance of core PBS proteins was insensitive to minor repression but decreased significantly as repression increased (Fig. 4F); this effect is consistent with our spectroscopic analysis (Fig. 4D). Iron-handling proteins exhibited a range of enrichment at low repression levels (i.e., Log_2_FC = -2 to 3), but decreased as repression increased (Fig. 4G). This reduction may reflect the role of iron in generating ROS, which could increase with NDH-1 depletion; in general, iron is tightly regulated in cyanobacteria (too much causes ROS, while too little limits photosynthesis) (42, 43). Not all proteins showed downward trends. Ribosomal proteins exhibited similar levels of depletion at all repression levels (Fig. 4H), and many proteins became more abundant as *ndhG* repression increased (Fig. 4I). Importantly, an increase in the abundance of DNA repair and high-light inducible proteins (e.g., SbcD and A0186) suggests that under 22B, minor repression of *ndhG* might elicit a general stress response that becomes more consequential—that is, more deleterious—as repression increases. Correlated enrichment trends for unannotated proteins, in turn, suggest that they may contribute to stress responses, though further characterization is required. In general, under 22W, all aforementioned trends were muted or absent (S14).

The benefits conferred by minor repression of *ndhG* under 22B and high repression under 22R are perplexing. Under both conditions, repression of this gene should decrease CET and temper NDH-1-mediated reduction of the plastoquinone (PQ) pool. Using fluorescence relaxation kinetics, we measured the impact of *ndhG* repression on the reoxidation of PSII-bound primary plastoquinone (Q_A-_) in cells grown under 22B and 22R. Our results showed subtle differences in reactivation kinetics between growth conditions: Under 22B, cells exhibited a small redox “wave” (SI Appendix, Fig. S9B). Prior work indicates that this wave results from transient oxidation of a highly reduced PQ pool by PSI, followed by re-reduction via NDH-1 (44); an insensitivity to *ndhG* reduction in our measurements, however, suggests that other photosynthetic machinery overexpressed in blue light (e.g., PetD, a subunit of cytochrome b6f; Fig. 4E) might play a role. At high *ndhG* repression, cells grown under 22B, but not 22R, exhibited a slow middle phase (SI Appendix, Fig. S9F)—an indication of slow reoxidation of bound plastiquinone (Q_A_^-^) at vacant sites of PSII centers, where mobile plastiquinone (Q_B_) arrives via diffusion (45)—likely as a result of cellular stress (e.g., as in Fig. 4D).

### Multi-gene transcriptional adjustments exhibit nonlinear effects

We used a dual-guide system to explore the additivity of transcriptional effects. In brief, we placed two guides on either side of an *aceEF* RNA hairpin, which is cleaved by RNase III inside the cell (Fig. 5A) (46). Dual repression of *ftsZ* and *cpcB* with this system produced expected cell elongation and pigmentation patterns, respectively, regardless of sgRNA order (SI Appendix, Fig. S10). We focused our analysis on genes that showed pronounced sgRNA enrichment in 22R or 22B (Fig. 5B). For each two-guide combination, we used maximally enriched guides in both orientations (i.e., AB and BA, where A and B are separate guides), which allowed us to capture RNA processing effects, which can increase the relative abundance of the first guide (47).

**Figure 5.**
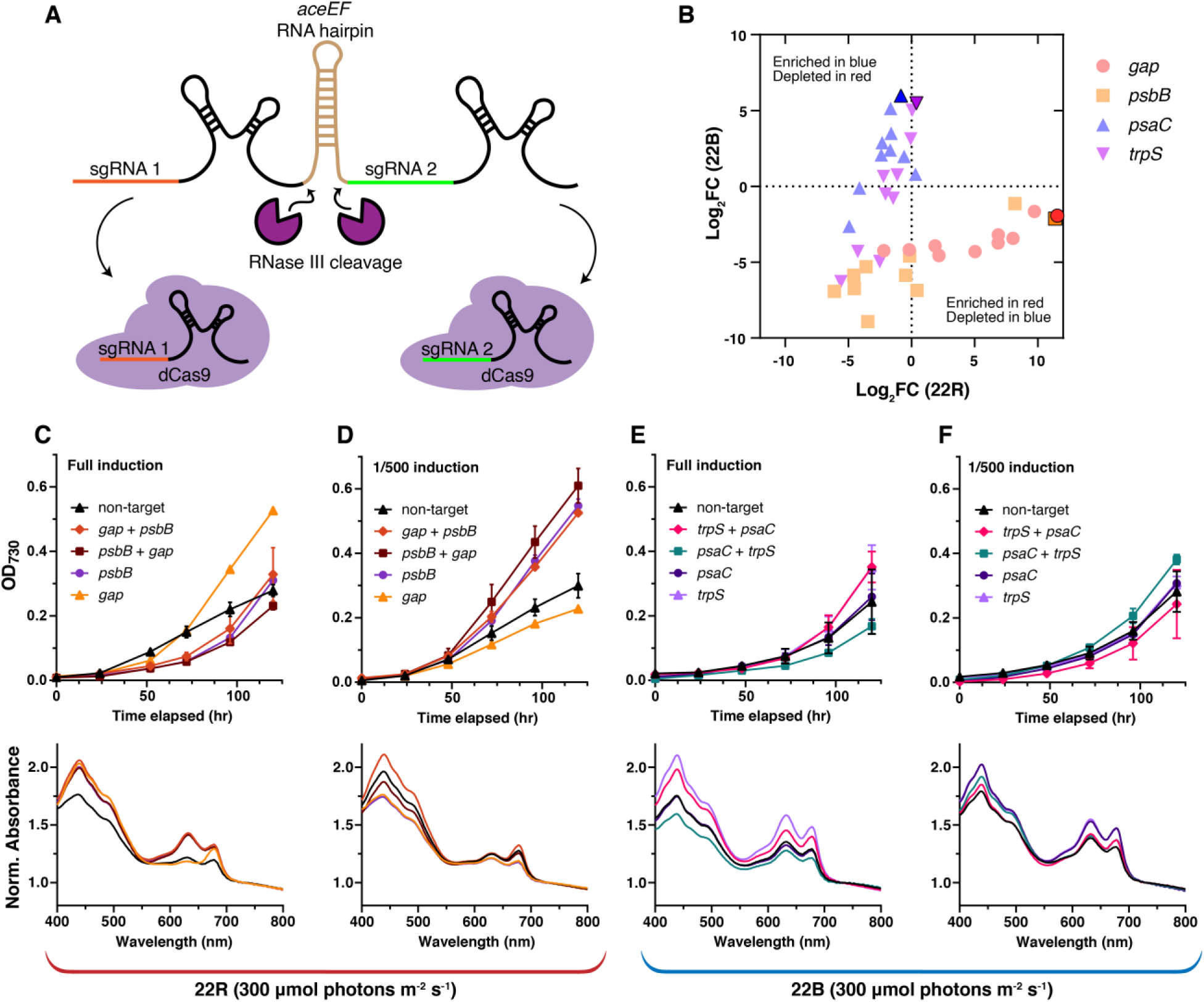
Dual knockdowns display non-linear effects. (A) Schematic of a single-transcript dual-sgRNA system. Native RNase III cleaves the *aceEF* RNA hairpin to generate two sgRNAs which complex with dCas9 (46). (B) The Log_2_FC of sgRNAs for genes that exhibited prominent sgRNA enrichment under 22R or 22B. Highlight: maximally enriched guides (black). (C-F), Growth curves and absorbance scans of single and double constructs grown in (C-D) red or (E-F) blue light at 300 µmol photons m^−2^ s^−1^ at different inducer concentrations: full (5 mM IPTG, 0.5 µg/ml aTc; C,E) or 1/500 (10 µM IPTG, 1 ng/ml aTc; D,F). Absorbance spectra are normalized to 730 nm. Growth curves evidence sensitivity to guide order and inducer concentration. Data represent the mean and standard deviation of n = 2 biological replicates.

Combinatorial transcriptional effects were nonlinear. Under 22R, the growth rate afforded by repressing *psbB* and *gap* was slower than repressing *gap* alone, which had a dramatic effect (Fig. 5C). The *psbB* and *gap* genes encode a PSII reaction center and a glyceraldehyde-3-phosphate dehydrogenase, respectively. Absorbance spectra show that *gap* repression reduces PBS content (Fig. 5C, 630 nm), which may alleviate an energy sink, as PBS accounts for up to 50% of the soluble protein in cyanobacteria (48). When we reduced the inducer concentration by 500-fold, in turn, the relative growth rates afforded by different guides flipped; repression of *psbB*—alone or in combination—produced the highest growth rates, while *gap* repression slowed growth below a non-target guide (Fig. 5D). PBS content remained similarly low in all cases.

Under 22B, transcriptional effects were subtle but showed extreme sensitivity to guide levels. Dual repression of *trpS* and *psaC*, which encode a tryptophanyl-tRNA synthetase and a PSI iron-sulfur center, produced both the fastest and the slowest growth rates, depending on guide order; and at low inducer concentrations, the effects on growth flipped (Figs. 5E and 5F). The influence of guide order and induction level on growth rates echoes the enrichment gradients observed in our genome-wide screen and shows how the precise level of transcriptional repression determines fitness.

## DISCUSSION

Cells adapt to changing environments by using biochemical circuits to regulate gene expression, often by degree. In this study, we used a high-density genome-wide CRISPRi screen to assess the influence of intermediate levels of transcriptional repression on growth under environmental extrema. Under cold monochromatic conditions, many knockdowns improved fitness, though these effects changed with the extent of repression—a function of guide strength, array organization, and induction level. The *ndhG* gene is illustrative: Minor repression improved growth under 22B, while high repression became toxic; under 22R, in turn, high repression improved growth. The NDH-1 complex plays an essential role in photosynthesis, CO_2_ uptake, oxidative stress, and respiration; the link between its impeded assembly and improved fitness under monochromatic extrema is intriguing and merits focused examination. Our ability to detect this influence, however, highlights the biochemical resolution afforded by sampling different levels of transcriptional repression in genome-wide CRISPRi screens and cautions against the use of binary classifiers of gene-specific essentiality.

The influences of low temperature and monochromatic light were nonadditive; most fitness-improving knockdowns emerged only when these conditions were combined. Under blue light, for example, cells increased the production of PSII and PSII-PBS, likely to address a photosystem imbalance; poor absorption by PBSs, however, makes this a (largely) ineffective strategy. In general, genes with fitness-improving knockdowns under red and blue light were distinct, evidencing unique stress responses and alleviation mechanisms. For red light, most of these genes related to energy metabolism; for blue light, the largest enriched fraction contributed to genetic information processing. The mechanisms by which transcriptional repression improves fitness are challenging to generalize, though our analyses highlight (i) amplification of a native cellular response (e.g., PSII assembly under blue light) and (ii) attenuation of an unnecessary resource (e.g., PBSs under red light) as strategies. Overall, our findings show how minor transcriptional adjustments can lead to major improvements in growth under environmental extrema and provide guidance (i.e., gene targets) for engineering well-adapted strains.

Regulatory pathways typically modulate the transcription of multiple genes. Our combinatorial experiments illustrate the complexity of two-gene adjustments, where fitness effects under environmental extrema were nonadditive, relative to single-gene changes, and showed sensitivity to both sgRNA arrangement (i.e., sgRNA order in an array) and induction level. These findings highlight the importance of genetic context and, for future work, motivate the use of large-scale dual-guide screens to uncover unexpected synergies or, perhaps, to generate functional assignments for unannotated genes based on the fitness effects exhibited by their joint repression with well-characterized genes. Broadly, CRISPRi screens carried out with high-density single- and dual-guide libraries across growth conditions can supply rich datasets for modeling the complex interplay of transcriptional regulation and environmental adaptation. The set supplied by this study already provides a comprehensive starting point for such efforts.

## MATERIALS AND METHODS

SI Appendix, SI Methods details methods for cloning. strain assembly, and sgRNA library design; screening procedures and analyses; and all biophysical experiments, including proteomics, 77K spectroscopy, and kinetic fluorimetry.

## AUTHOR CONTRIBUTIONS

A.H., J.C.C., J.M.F., and C.E. conceived of research, designed experiments, and analyzed data. A.H. and N.L. carried out sgRNA design, growth experiments, NGS analyses, spectroscopic studies, and data processing. D.L.C. and P.A. performed proteomics analyses. A.H., J.M.F., and C.E. wrote the manuscript with comments from all authors.

## ACKNOWLEDGEMENTS

This work was supported by the U.S. Department of Energy, Office of Science, Office of Biological and Environmental Research, under Award Number DE-SC0018368 (A.H., D.L.C., P.A., J.C.C., and C.E.), and the United States Army Research Office, under Award Number W911NF-18-1-0159 (J.M.F.). The work (proposal:508377) conducted by the U.S. Department of Energy Joint Genome Institute (https://ror.org/04xm1d337), a DOE Office of Science User Facility, is supported by the Office of Science of the U.S. Department of Energy operated under Contract No. DE-AC02-05CH11231.

