## Supplementary Information for "High-density CRISPRi screens reveal adaptive transcriptional gradients in cyanobacteria"

SI Methods. . . . . . . . . .. . . . . . . . . . . . . . . . . . . . . . . . . . . . . . . . . . . . . . S2

Figures S1-S14. . . . . . . .. . . . . . . . . . . . . . . . . . . . . . . . . . . . . . . . . . . . S13

Tables S1-S7 . . . . . . . . . . . . . . . . . . . . . . . . . . . . . . . . . . . . . . . . . . . . . S29

SI References . . . . . . . . . . . . . . . . . . . . . . . . . . . . . . . . . . . . . . . . . . . . . S36

### SI METHODS

**Materials.** We purchased chemicals from Thermo Fisher Scientific (NaCl, $MgSO_4$*7$H_2O$, KCl, $CaCl_2$*2$H_2O$, $NaNO_3$, $KH_2PO_4$, Tris HCl, IPTG, DMSO) MP Biomedicals (TAE), IBI Scientific (agarose), Teknova (LB broth, LB agar, agar), chemodex (anhydrotetracycline; aTc), and MilliporeSigma (kanamycin). We purchased restriction enzymes, Q5 polymerase, and T4 DNA ligase and kinase from New England Biolabs.

***E. coli* strains.** We used chemically competent NEB DH5a (#C2987H) and Zymo Mix and Go! DH5 Alpha (#T3007) cells for cloning and Lucigen E. cloni 10G (#600521) electrocompetent cells for library preparation.

**Cloning and molecular biology.** We constructed plasmids with Gibson and Golden Gate assembly. Table S3 describes the composition of each plasmid; Table S4 lists primers used for plasmid assembly and NGS sequencing.

**Design of sgRNA libraries.** To build a high-density library, we selected ten sgRNAs for each annotated gene in the genome of *Synechococcus* sp. PCC 7002 (NCBI accession ASM1948v1) by using a custom python script (available at https://github.com/aphren/Hren-2024-scripts). Briefly, we calculated a “seed score” for each possible guide by calculating the length of the shortest unique PAM-proximal sequence, relative to every possible NGG and NAG target site within the genome (Fig. S1), and we used this score to calculate the final rank score by incorporating distance from the annotated start codon as shown in Equation 1.

$$\text{rank score} = \frac{\text{distance from start codon (bp)}}{250\text{ bp}} + \text{seed score} \qquad (1)$$

For each gene, we selected the top ten sgRNAs (as indicated by low rank score) that bind to the coding strand. For genes with fewer than ten coding strand-targeting sgRNAs, we added sgRNAs targeting the noncoding strand. A handful of sgRNAs target genes with duplicates elsewhere in the genome and are annotated with all target loci. As controls for normalization, we included 780 non-targeting sgRNAs in the final library by randomly generating sequences and selecting guides with low seed scores (seed score of 7 for 98.7% of non-targeting guides). The library was synthesized by the Joint Genome Institute (JGI) under proposal #508377.

**Library preparation.** Our final sgRNA contained 32,928 unique sequences. To enables library-wide screens, we used a high-efficiency transformation protocol developed in our recent work**.** For review, we modified a previous method (1) by altering the sgRNA integration site, the length of its homology arms, and the cell density during transformation; the final protocol affords an efficiency similar to that of chemically competent E. coli (Fig. S4). A detailed description follows: To begin, we picked a colony of scJC0010 cells (Δ*acsA* dCas9 F RBS aTc; pCas2F (2)) to inoculate 25 mL liquid A+ media (3), which we grew at 37°C under 150 μmol photons $m^{-2}$ $s^{-1}$ of white light in ambient air and 200 rpm—hereafter, "standard conditions"—to an $OD_{730}$ of ~0.3 (Thermo Scientific Genesys 10S Vis). Here, our A+ media consisted of 18 g/L NaCl, 5 g/L $MgSO_4$*7$H_2O$, 30 mg/L $Na_2EDTA$*2$H_2O$, 600 mg/L KCl, 370 mg/L $CaCl_2$, 1 g/L $NaNO_3$, 50 mg/L $KH_2PO_4$, 1 g/L Tris HCl, 4 μg/L cyanocobalamin and trace metals at a pH of 8.2. From this culture, we inoculated a 25 ml culture of A+ media at $OD_{730}$ of 0.01 and grew the new culture under standard conditions to an $OD_{730}$ of ~0.3. We concentrated the cells via centrifugation (4,300 x*g*, 10 min) to an $OD_{730}$ of 6.0, mixed in the plasmid library, and incubated the cells in a capped microcentrifuge tube for 2 hours at 30°C under 60 μmol photons $m^{-2}$ $s^{-1}$.

We spread the transformations on ten 150 mm plates (A+ media, 50 µg ml$^{-1}$ kanamycin, and 1% agar) using glass beads and incubated the plates in standard conditions with no shaking for 3 days. We scraped over one million colonies from the plates, resuspended them in liquid A+ media (50 µg ml$^{-1}$ kanamycin) and grew the cell mixture in standard conditions for 2 hours. We concentrated the culture to an $OD_{730}$ of 3.0 in liquid A+ media with DMSO (10%; v/v) via centrifugation, and stored the cells in aliquots at -80°C. We repeated the process independently for the second biological library replicate.

**Strain construction.** We prepared cells with genomically integrated copies of dCas9 and a sgRNA under the control of inducible promoters by adapting a protocol from Moore *et al.* (1). For single guide constructs, we designed two oligos with an sgRNA spacer or its reverse complement and overhangs matching the pAPH07 insertion site, combined and phosphorylated them using T4 PNK (NEB), and performed a 15 min gradient heat inactivation and annealing step (95ºC start, -2ºC/min). We digested pAPH07 plasmid with BspQI at 50°C for 5 hours and gel extracted the construct to reduce background DNA. We used T4 DNA ligase to insert the phosphorylated spacer fragments into digested pAPH07 backbone; this reaction used a 7:1 molar ratio (insert : backbone) and overnight incubation at room temperature. After heat inactivation of the ligase (i.e., 98ºC for 5 min), we transformed NEB DH5a cells with the ligation product and grew them on selective plates (25 g/L LB, 50 µg ml$^{-1}$ kanamycin, and 1.5% agar). We picked colonies to inoculate 5 ml LB media (25 g/L LB with 50 µg ml$^{-1}$ kanamycin) for overnight growth at 37°C and 200 rpm, extracted the plasmids (Qiagen QIAprep Spin Miniprep Kit), and sequenced them to confirm the presence of the spacer. We transformed strain scJC0010 with the plasmid using natural transformation and grew transformants on agar plates (A+ media, 50 µg ml$^{-1}$ kanamycin, and 1% agar) under standard conditions with no shaking. For double guide constructs, we mixed oligos with pAPH021-compatible overhangs and phosphorylated as above, then inserted them via Golden Gate into pAPH021 (PaqCI-digested and gel extracted) alongside a PCR-amplified central insert (i.e., the region between the two sgRNA spacers). We incubated reactions for 50 cycles (37ºC, 5 min; 16ºC, 5 min), heat inactivated, then transformed as above.

**Library screens.** For each screening condition, we began culture preparation by filling two 2.8 L Fernbach flasks with 300 ml liquid A+ media (50 µg ml$^{-1}$ kanamycin). We thawed frozen library stocks on ice, vortexed them lightly to mix, and inoculated 500 µl into each replicate flask. We grew flasks in standard conditions for two hours, added inducer to 5 mM IPTG and 0.5 µg ml$^{-1}$ aTc, returned the flasks to standard conditions for two hours, then adjusted the incubator to the appropriate temperature and lighting conditions. We grew diurnal samples in a 12-hour light-dark cycle in standard conditions. For colored light, we positioned two LED lightbulbs (HIGROW 36W LED Plant Grow Light Bulb, 460 or 660 nm; Fig. S11) on opposite ends of the incubator to deliver a photosynthetic active radiation (PAR) intensity of 150 µmol photons m$^{-2}$ s$^{-1}$ to the center of each flask (LI-COR LI-250A Light Meter). At each timepoint, we (i) removed 500 µl of culture to measure $OD_{730}$ and (ii) switched flask positions to mitigate subtle differences in light intensity that might exist between different regions of the shaker. We grew cultures until both replicates exceeded an $OD_{730}$ of 0.3 (~9-10 population doublings; Fig. S12), and pelleted and froze the cells at -20ºC.

**Sequencing and data analysis.** We used next-generation sequencing (NGS) to examine changes in sgRNA populations between the start and end of each growth condition. To begin, we extracted genomic DNA (Zymo Quick-DNA Fungal/Bacterial Miniprep Kit) and used PCR to amplify the sgRNA coding sequence (Q5 2x Master Mix); for each reaction, we used at least 300 ng of gDNA template and mixed primers containing 0-3 randomized bases on the 5' end, which introduces phasing (4) (Table S4). We extracted (Qiagen QIAquick Gel Extraction Kit**,** Econospin All-In-One DNA Spin Columns) the amplicons from a 2% agarose gel using Promega X-tracta Gel Extractors to get ~30-100 ng of DNA (Qubit dsDNA HS Assay). We prepared samples for sequencing using Illumina's NEBNext Ultra II DNA Library Prep Kit and NEBNext Multiplex Oligos (96 Unique Dual Index Primer Pairs) and sent final samples for sequencing (NextSeq 2x150 bp Illumina at Azenta) to obtain a total of ~700M reads across all samples. We used VSEARCH (5) to process our sequencing data. Briefly, we merged raw paired-end reads using the "fastq_mergepairs" command (id = 1.0), converted merged files to FASTA format using "fastq_filter", aligned reads to our sgRNA database using "usearch_global" (id = 1.0), and tabulated final counts from the alignment output. We calculated the $log_2$-fold change ($Log_2FC$) enrichment by (i) normalizing read counts to the negative binomial distribution model of DESeq2 (6) and (ii) shrinking $Log_2FC$ values using the "apeglm" approximate posterior estimation method (7) to reduce variance while preserving large effect sizes. We identified genes with significant changes in sgRNA frequency using CRISPhieRmix (8). Collated DESeq2 output and read counts are listed in Table S1.

**Essentiality comparison:** We compared our results to the results of two other genome-wide screens by examining the fitness effects associated with homologous proteins. To begin, we identified protein homologs between PCC 7002 and either *Synechococcus elongatus* PCC 7942 or *Synechocystis* sp. PCC 6803 by using KBase's GenomeProteomeComparison (v0.0.8). For our comparisons with PCC 6803, we assigned a “significant” classifier to PCC 7002 genes with a non-null designation in at least one of seven conditions, as determined via CRISPhieRmix (< 0.1 local FDR). For our comparison with PCC 7942, we further filtered genes with a signficant classification in PCC 7002 by requiring a mean $Log_2FC$-change less than zero in at least one condition to assign a “depleted” classifier. We gathered assignments for significance in PCC 6803 (9) and essentiality in PCC 7942 (10) from literature. For PCC 7002 proteins with multiple homologs, if at least one homolog met our classification criteria in the corresponding cyanobacterium, we considered it to be depleted or significant.

**Gene clustering:** We used k-means clustering to group genes by shared sgRNA enrichment patterns. Our analysis used (i) the mean sgRNA enrichment for each gene and (ii) the number of enriched sgRNAs (i.e., $Log_2FC > 1$) for genes with one or more significantly enriched guides (padj < 0.05) to create five clusters. Here, we normalized the $Log_2FC$ enrichment data for 37W, 22B, and 22R conditions with the scikit-learn (11) StandardScaler and clustered the normalized data with the scikit-learn KMeans clustering algorithm. For visualization, we used the scikit-learn t-SNE function (elkan algorithm and n_init = 200) and matplotlib (12). The $Log_2FC$ means and cluster assignments are listed in Table S5.

**Gene ontology:** To compare the gene ontology in the five clusters depicted in Figure 3, we downloaded ontology data from Kegg (13) and extracted the ontology assignments for each gene. We identified the three most abundant ontology categories within each cluster and calculated their relative proportions. We quantified gene fraction as the proportion of genes in a cluster that belong to the three most abundant categories, relative to all genes within the same cluster.

**Digital PCR.** We used digital PCR to evaluate the influence of a select subset of sgRNAs for *ndhG* on transcription. In brief, we transformed cells with low-, medium, and high-strength guides (Table S7), used transformed strains to inoculate liquid A+ media (50 µg $ml^{-1}$ kanamycin, 5 mM IPTG, 0.5 µg $ml^{-1}$ aTc) in 125 ml Erlenmeyer flasks, and grew each culture under standard conditions to an $OD_{730}$ of 0.3 - 0.5. We mixed 5 ml from each final culture with 10 ml Qiagen RNAprotect Bacteria Reagent, pelleted the cells (4300 x*g*, 10 min), extracted total RNA using the Qiagen RNAeasy Mini Kit, and converted extract RNA to cDNA using the ThermoFisher SuperScript IV First-Strand Synthesis System. We ordered TaqMan dye-labeled probes complementary to *ndhG* (A0924; FAM dye) and *dnaK* (A2160; VIC dye) from ThermoFisher. We quantified *ndhG* transcript abundance relative to *dnaK* with the QuantStudio 3D Digital PCR System.

**Growth comparisons:** We compared the growth rates of isolated PCC 7002 strains containing different CRISPRi systems by growing them in A+ media (15-25 ml in 125 ml Erlenmeyer flasks) at 200 rpm under ambient air. We used a Percival AL-41L4 photoincubator with 6-8 LED bulbs (HIGROW 36W LED Plant Grow Light Bulb, 460 or 660 nm) arranged in a symmetric orientation over the cultures and calibrated the PAR intensity with a light meter (Gigahertz-Optik MSC15). For our analysis of different *ndhG* guide strengths or induction levels (i.e., Fig. 3D), we inoculated cultures from stock liquid culture into fresh A+ media with the requisite inducers (50 µg $ml^{-1}$ kanamycin, 5 mM IPTG, and 0.5 µg $ml^{-1}$ aTc) and grew them for three days at 150 μmol photons $m^{-2}$ $s^{-1}$ PAR. For pooled *ndhG* sgRNAs (i.e., Fig. 3E), we prepared freezer stocks by mixing stock cultures of each strain at an $OD_{730}$ of 0.3, pelleting the cells, resuspending them in A+ media with 10% DMSO (v/v), and freezing them at -80°C. For each light intensity (75 – 450 μmol photons $m^{-2}$ $s^{-1}$), we inoculated two flasks from a single freezer stock and grew them to an $OD_{730}$ of 0.2. We measured changes in sgRNA frequency via NGS (Quintara Biosciences AmpExpress). For comparisons of single- and double-guide constructs, we inoculated cultures from frozen aliquots, incubated for two hours in standard conditions, induced and incubated flasks for two more hours, and grew them for five days under 450 nm or 660 nm light (300 μmol photons $m^{-2}$ $s^{-1}$). For all experiments, we measured $OD_{730}$ with a Spectramax iD3 microplate reader.

**77 K fluorescence:** We carried out cold-temperature spectroscopy to analyze photosynthetic machinery. Briefly, we grew wild-type PCC 7002 cells in either 22W, 22R, or 22B to an $OD_{730}$ greater than 0.3, diluted these cells in A+ media to an $OD_{730}$ of 0.15, and returned them to the incubator for 30 minutes. We transferred each culture to 5 mm glass NMR tubes and rapidly froze them in liquid nitrogen. We performed 77 K fluorescence emission scans by exciting at 440 nm (chlorophyll *a*) or 580 (phycocyanin) on a Horiba Fluorolog FL-1000 FL3-11 (14). For normalization, peak wavelengths were chosen within acceptable ranges of those used in previous work (15, 16).

**Proteomics.** We used quantitative proteomics to examine changes in protein abundance caused by different levels of *ndhG* repression. To begin, we selected three sgRNAs targeting *ndhG* (Table S7) based on their differential fitness effects in 22B (negative, neutral, and positive) and cloned them into pAPH07. We transformed these constructs, alongside a non-targeting sgRNA construct (nt-gRNA_11), into scJC0010 and grew the transformed cells on agar plates (A+ media, 50 µg $ml^{-1}$ kanamycin, and 1% agar). We picked colonies from each plate to inoculate four 15 ml liquid cultures (A+ media with 50 µg $ml^{-1}$ kanamycin), which we grew at 37°C and 200 rpm under 180 µmol photons $m^{-2}$ $s^{-1}$ white light for 136 hours. We concentrated the cultures 10-fold (15ml to 1.5ml) into liquid A+ media with 10% DMSO (v/v) and stored them at -80ºC.

We completed growth experiments in 22B and 22W as in the library screen. In brief, we thawed each sgRNA-specific freezer stock on ice, vortexed it briefly, and used 40 µl to inoculate liquid A+ media (50 µg $ml^{-1}$ kanamycin) in 125ml Erlenmeyer flasks with foam stoppers. We grew cells at 37°C and 200 rpm under 180 µmol photons $m^{-2}$ $s^{-1}$ for two hours, added inducer and returned flasks to the incubator for two hours, then transferred cultures to their respective conditions (i.e., 22B or 22W). Each day, we scrambled the cultures to reduce the impact of uneven light exposure and tracked $OD_{730}$ by removing 200 µl culture samples (Tecan Spark microplate reader).

For each sample, we processed cellular biomass pelleted by centrifugation (4,300 x*g*, 10 min) for liquid chromatography tandem mass spectrometer-based proteomic measurements. We performed cellular lysis, protein precipitation, and protease digestion as previously described (17). In brief, we lysed cells via bead beating and a detergent-based lysis buffer. We precipitated and isolated proteins using the protein aggregation capture (PAC) method (18) and digested them in situ with sequencing-grade trypsin. We analyzed peptides by automated 2D-LC-MS/MS and processed them as previously described (19). Briefly, we separated peptides on an analytical column (75 µm x 350 mm) packed with Kinetex RP C18 resin and analyzed samples using a Top10 data dependent acquisition strategy (20). We analyzed resulting LC-MS/MS data using the Proteome Discoverer software (Thermo-Fisher Scientific, Version 2.5). For the identified proteins, we performed Log2-transformation of abundances followed by local regression (LOESS) normalization and mean-centering across the entire dataset in R using scripts from the InfernoRDN software(21). We imputed the abundance values for proteins with missing values with random values drawn from the normal distribution (width .3, downshift 2.2) using R. We used NMDS statistical analysis to confirm there was strong agreement between replicates (Fig. S13). We deposited all proteomics spectral data in this study at the ProteomeXchange Consortium via the MASSIVE repository (https://massive.ucsd.edu/). The data can be reviewed under the username "reviewer_ MSV000094564" and password "Cyano". The processed proteomics data and statistical analysis are listed in Table S2.

**Design of a dual-guide system.** We designed a dual-guide system compatible with short-read sequencing of both sgRNA spacers (Illumina; paired-end sequencing). Guided by previous designs for post-transcriptional modification (22), we separated two sgRNAs with a well-studied RNA hairpin (*aceEF*) (23), which is cleaved by RNAse III, and evaluated its ability to create two functional sgRNAs in PCC 7002. Here, we selected two gene targets, *ftsZ* and *cpcB*, where transcriptional repression yields visual phenotypes (cell elongation and altered pigmentation, respectively). We used a Nikon TiE microscope and the supplied software (Nikon NIS-Elements AR, version 5.11.00, 64-bit) with a 100x oil immersion objective (Nikon, CF160 Plan Apochromat, NA = 1.45) for imaging. We imaged chlorophyll and phycobilin fluorescence using the Cy5 (645 nm peak excitation, 705 nm peak emission, 2% laser power of 0.368 mW) and RFP (555 nm peak excitation, 665 nm peak emission, 48% laser power of 18.2 mW) filter sets, respectively. We adjusted images to the same brightness threshold using Fiji (24).

**Kinetic fluorimetry:** We used kinetic fluorimetry to measure levels of PSII-bound PQ in cells. In brief, we grew cells in liquid A+ media (15 ml in 125 ml Erlenmeyer flasks) at 22°C and 200 rpm under 300 µmol photons $m^{-2}$ $s^{-1}$ of blue or red light and ambient air. We diluted the cultures to an $OD_{730}$ of 0.3 and analyzed samples using a fluorimeter (Photon Systems Instruments Double-Modulation Fluorometer FL 6000). We dark-adapted samples for >5 minutes, then took measurements using a $Q_A$- reoxidation protocol and fit the data to Equation 2, a standard multicomponent kinetic equation (25). Here, F(t) – $F_0$ is relative fluorescence yield, $A_{1\text{-}3}$ are decay amplitudes, and $T_{1\text{-}3}$ are time constants.

$$F(t) - F_0 = A_1 \exp\left(-\frac{t}{T_1}\right) + A_2 \exp\left(-\frac{t}{T_2}\right) + A_3/(1 + \frac{t}{T_3}) \qquad (2)$$

**Data and materials availability**: All data are available in the main text or the supplementary materials.

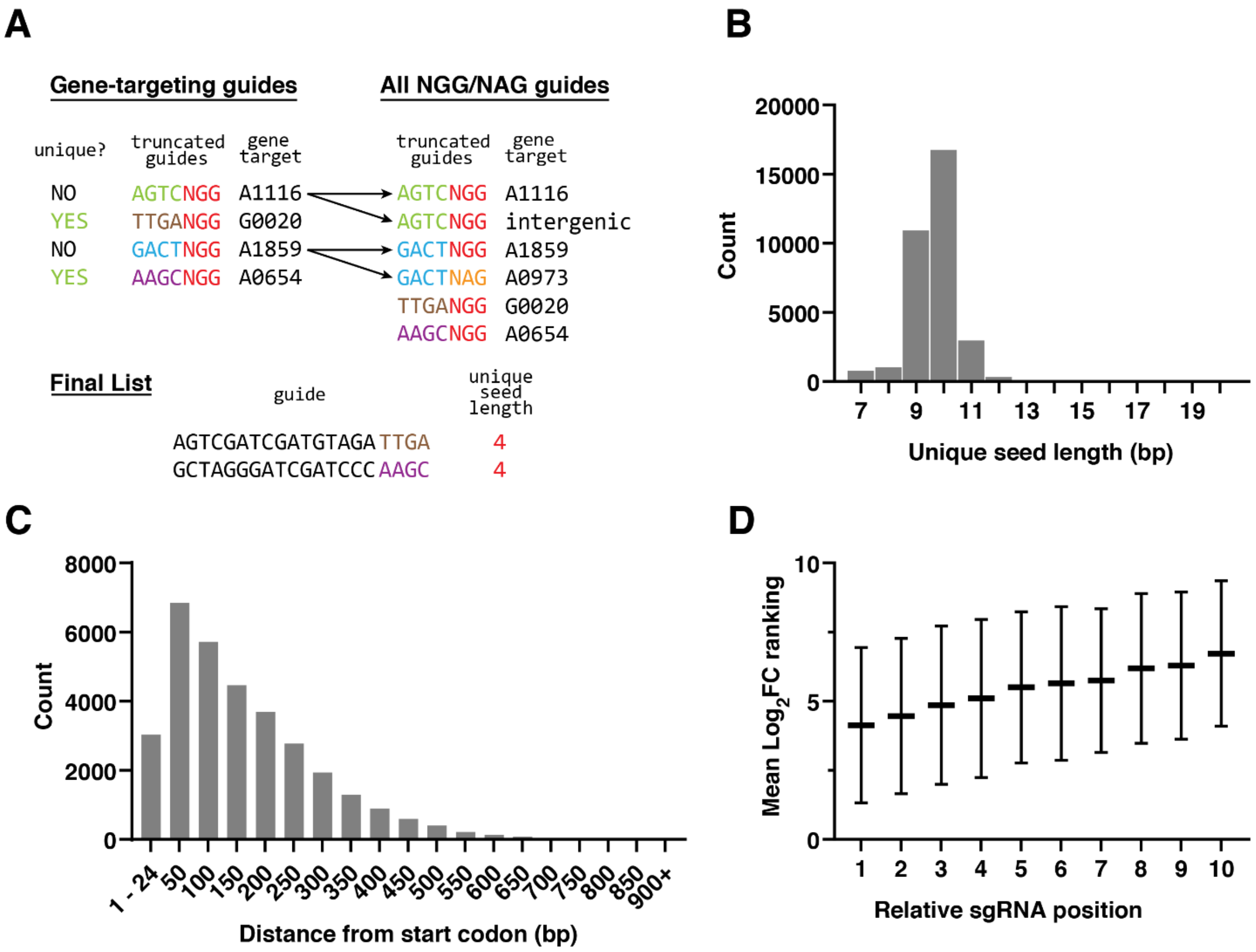


**Fig. S1. Design of sgRNAs for a genome-wide study of transcriptional repression in PCC 7002.** (A) For each possible sgRNA, our software package determines the length of the unique seed sequence in iterative steps: (i) It truncates the sgRNA sequence at the 3’ end and compares it to all possible NGG/NAG-containing genomic target sequences truncated to the same length. (ii) If the truncated sequence is not unique, the software adds an additional base and repeats step i. (iii) If the sequence is unique, it assigns a score equal to the minimum unique truncation. (B) For our study, the majority of sgRNAs had a unique seed length of 9-10. That is, they do not have a unique eight-base seed, but the ninth or tenth base differentiates them from all other possible seed sequences of the same length. (C) Our software package prioritizes seed sequences proximal to the start codon of the coding strand. Displayed distances represent the upper limit for each bin, containing all distances greater than the previous bin max. The majority of seed sequences sit within the first 200-300 bp of the start codon. (D) We examined the influence of binding position within a gene—here, “relative sgRNA position”—on repression strength by examining all sgRNAs with a $Log_2FC$ less than -1 in standard growth conditions (37W). For each gene, we ranked both (i) sgRNAs by their distance from the start codon (1 = closest and 10 = furthest away) and (ii) $Log_2FC$ values by their magnitude (1 = largest and 10 = smallest). This plot shows average sgRNA rank for each position rank across all genes. Errors bars represent the standard deviation in $Log_2FC$ ranking. Consistent with prior CRISPRi studies, proximity to the start codon correlates loosely with $Log_2FC$.

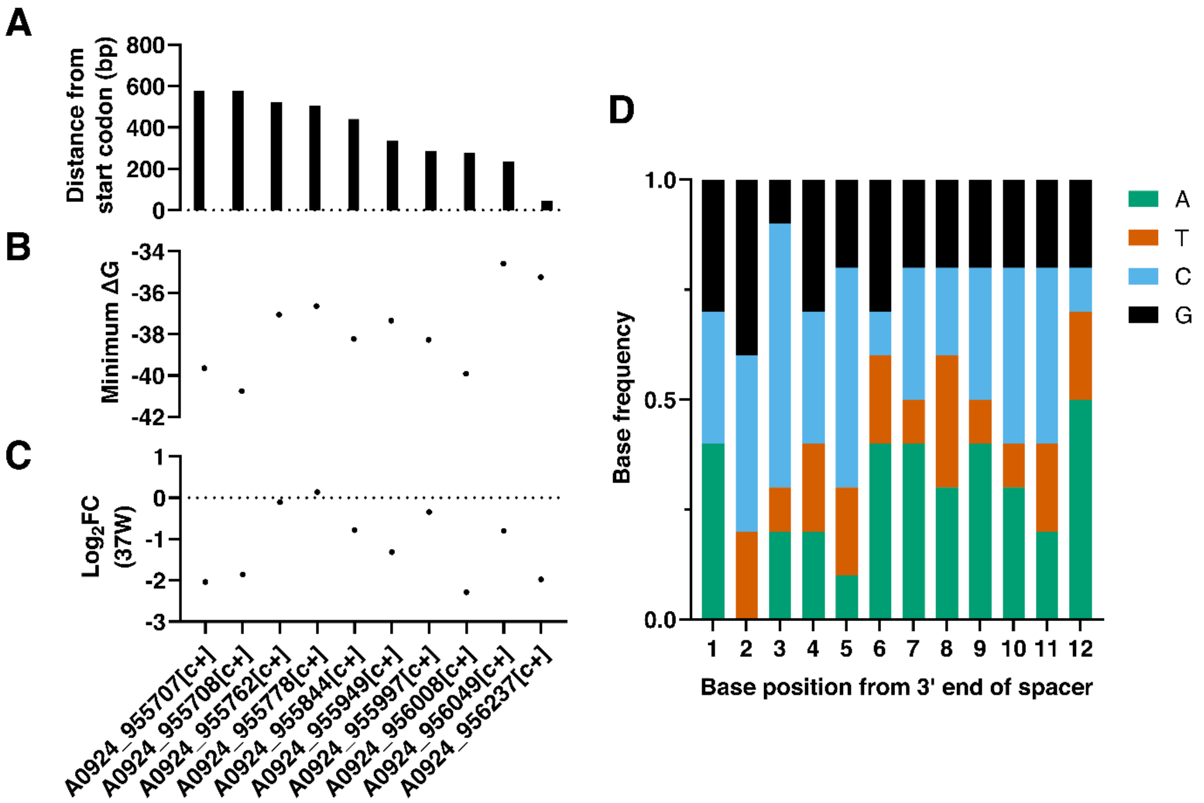


**Fig. S2. Guides targeting the same gene have diverse properties.** For the ten guides designed to target *ndhG*, there is large variability in the binding distance from (A) the start codon, (B) the minimum ΔG, and (C) the $Log_2FC$ in 37W. The composition of seed sequences exhibits (D) natural variability. Minimum ΔG of the full-length sgRNA was calculated using the IDT OligoAnalyzer hairpin prediction tool.

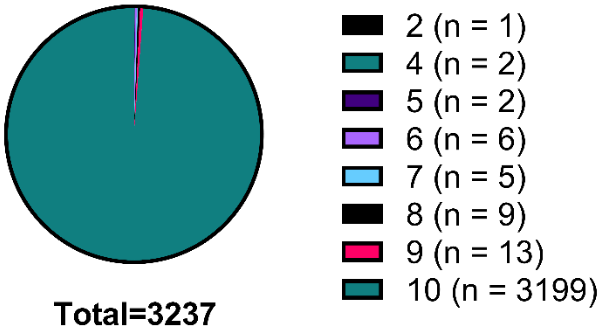


**Fig. S3. A high-density library of sgRNAs.** We designed an average of 10 sgRNAs per gene. The final library contains fewer than 10 sgRNAs for a small number of genes (38) due to their short length or a dearth of unique target sites. In the end, 3199 genes had 10 guides, constituting 98.8% of the library.

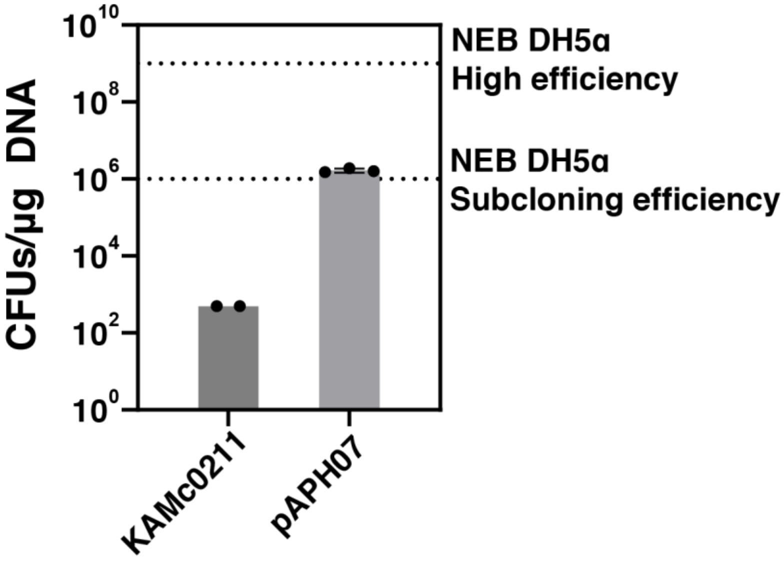


**Fig. S4. Transformation efficiency on par with commercial chemically competent *E. coli*.** In prior work, we developed a high-efficiency transformation protocol, the results of which are shown here in comparison to the basis vector and protocol (1). In brief, we transformed the scJC0010 strain with one microgram of KAMc0211 or pAPH07 plasmid using the original or updated transformation protocol, respectively. We performed spot plating to calculate the efficiency of transformation into PCC 7002, which exceeded the minimum efficiency advertised by New England Biolabs for their subcloning grade DH5α competent *E. coli* (Cat. #C2988J). Data depict n≥2 technical replicates.

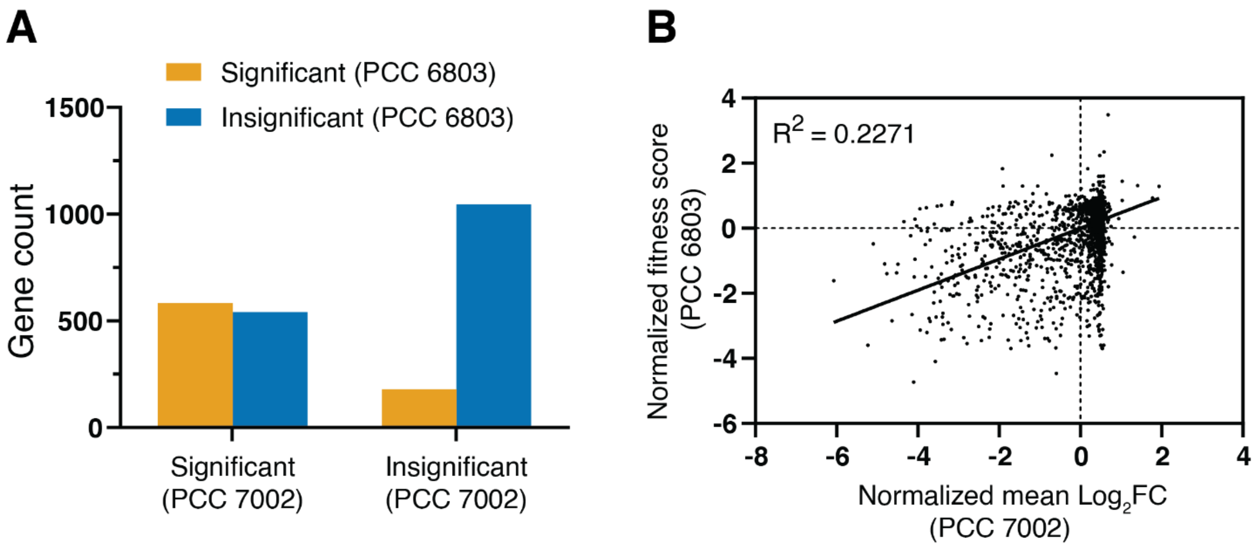


**Fig. S5. Pooled CRISPRi in *Synechocystis* sp. PCC 6803 shows strong gene overlap with PCC 7002.** (A) Proteins with significant fitness scores in a genome-scale CRISPRi screen in *Synechocystis* sp. PCC 6803 (9) showed strong overlap with protein homologs in PCC 7002. (B) No clear correlation exists between the fitness scores calculated in previous work (9) and the mean fold change in PCC 7002.

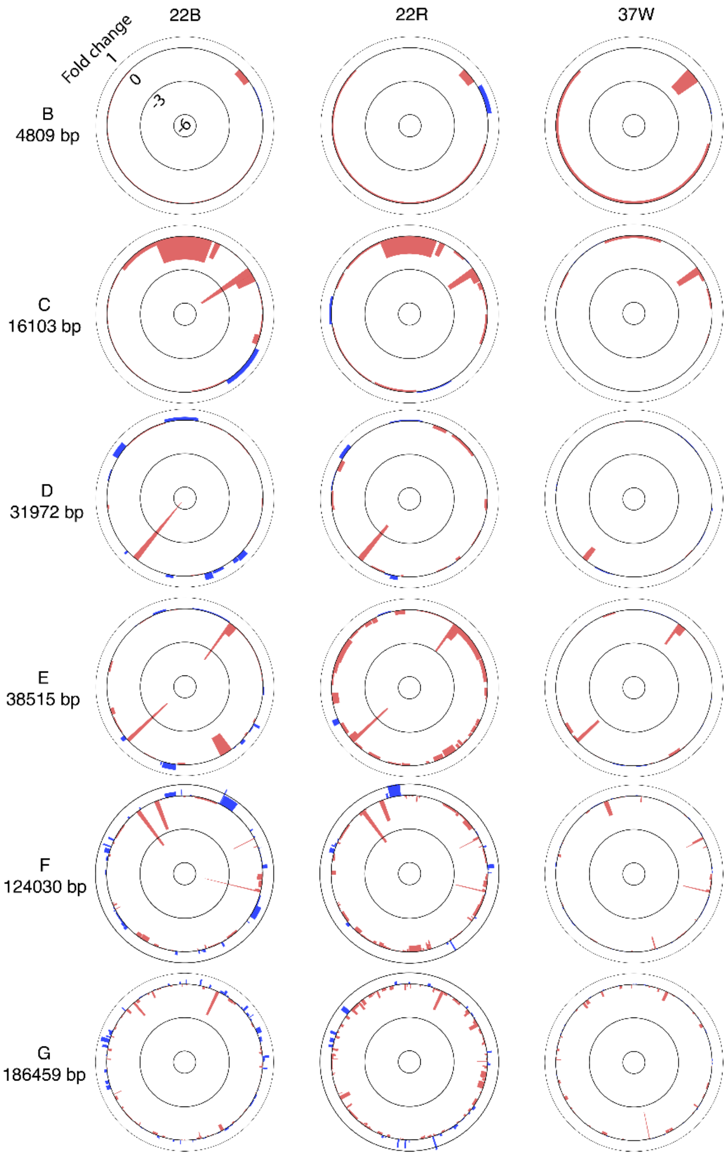


**Fig. S6. CRISPRi analysis of endogenous plasmids in PCC 7002.** We examined the mean sgRNA enrichment for plasmid-borne genes caused by growth in 22B (column 1), 22R (column 2), and 37W (column 3). Bar positioning matches gene location, and bar width matches gene size relative to plasmid size. The plasmids are arranged from smallest (plasmid B) to largest (plasmid G), top to bottom. When compared to the genome, plasmids show similar but less dramatic trends in sgRNA enrichment—an effect that may reflect the buffering effects of plasmid copy number or the lower frequency of important genes on endogenous plasmids.

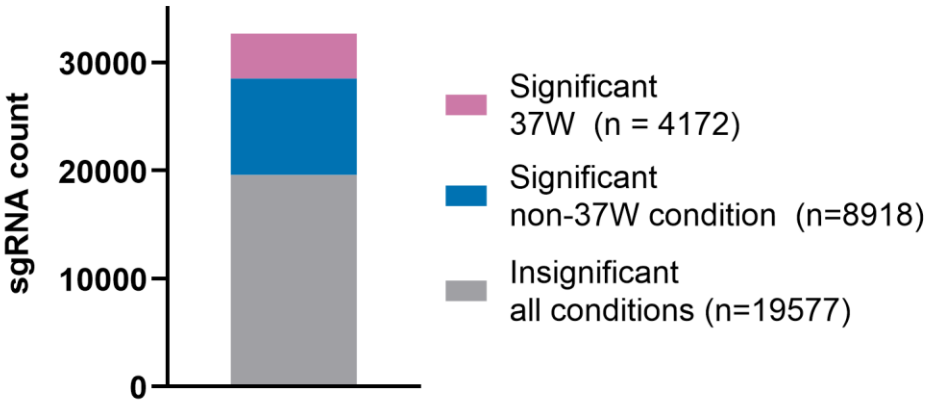


**Fig. S7. Guide depletion is condition dependent**. We categorized sgRNAs based on DESeq2-calculated adjusted p-values (padj). Only 12.8% of guides had a statistically significant change (i.e., depletion or enrichment) in standard growth conditions (37W, padj < 0.05). Of the remaining guides with insignificant effects in standard conditions, 31.3% induced a statistically significant fold change in at least one other condition – a population of sgRNAs that would appear ineffective without testing in environmental extrema.

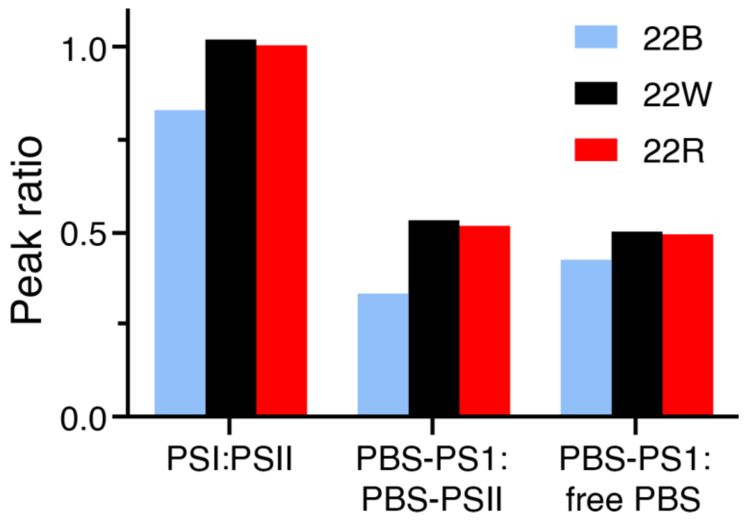


**Fig. S8. Cells grown in cold blue exhibit reduced 77 K fluorescence ratios.** We used normalized fluorescence values to calculate peak ratios (PSI:PSII; 720:685 nm, PBS-PS1:PBS-PSII; 712:685 nm, PBS-PS1:free PBS; 712:660 nm) from fluorescence emission spectra (Figs. 4A and 4B in the main text). Cells grown in cold blue exhibited reduced PS1 relative to PSII, as well as reduced PBS-PSI relative to both free PBS and PBS-PSII complex.

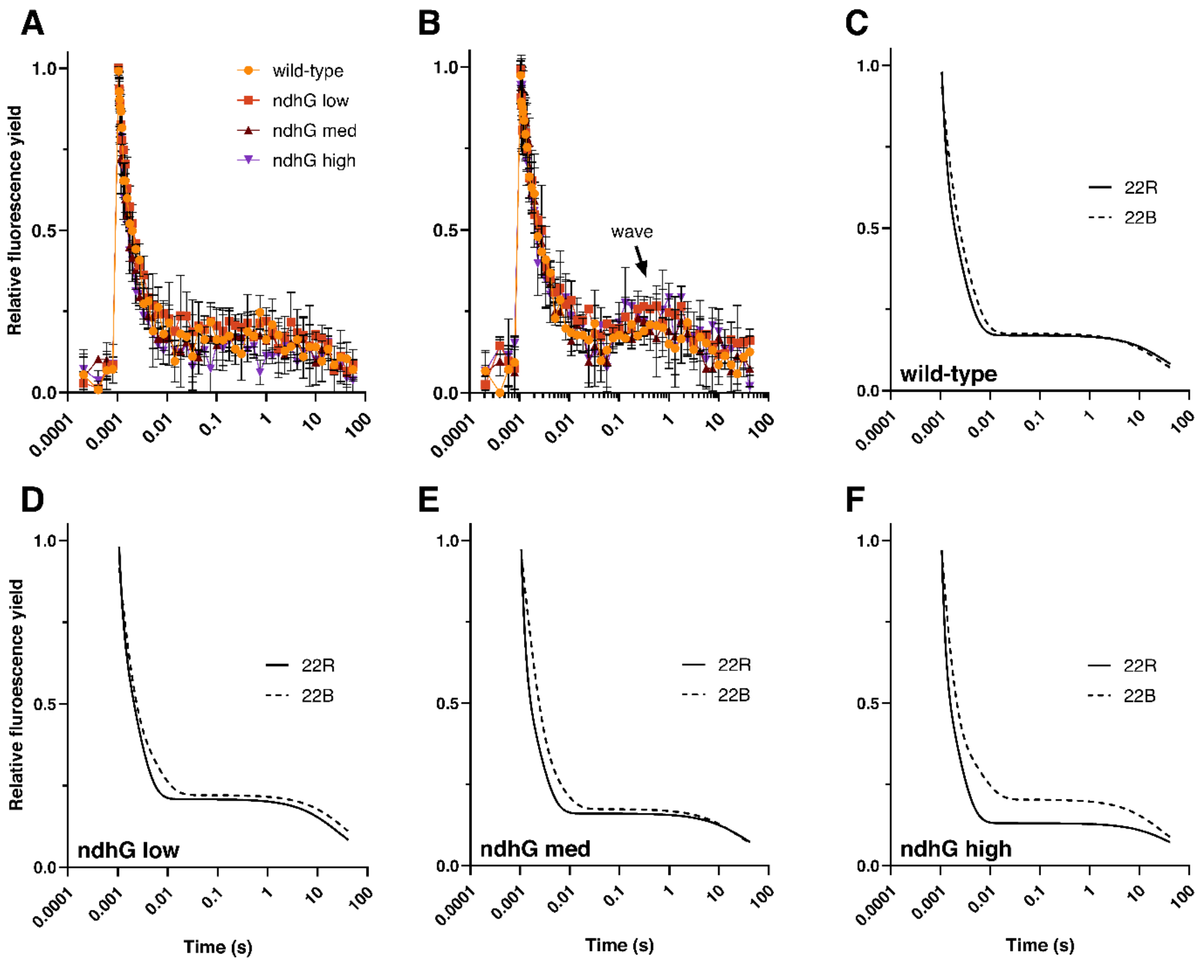


**Fig. S9. $Q_A$- reoxidation kinetics in cold red and blue.** (A-B) We grew cells under 300 μmol photons $m^{-2}$ $s^{-1}$ of (A) red or (B) blue light at 22°C, then measured the fluorescence relaxation kinetics of OD-normalized cultures following a single actinic flash of light. We normalized data relative to the minimum and maximum observed fluorescence within each time series. We observed a wave feature in cells grown in 22B, with fluorescence yield increasing briefly during the middle phase (i.e., 0.06 to 0.6 s). (C-F), We fit normalized data to a multicomponent kinetic decay equation for $Q_A$- reoxidation (fast phase – exponential, middle – exponential, slow – hyperbolic), then plotted curves across conditions for (C) wild-type, (D) *ndhG* low, (E) *ndhG* med, and (F) *ndhG* high samples. The wave feature is precluded in curve fitting due to the absence of a representative term in the decay equation. Data (A-B) depict the mean and standard deviation for n=3 measurements.

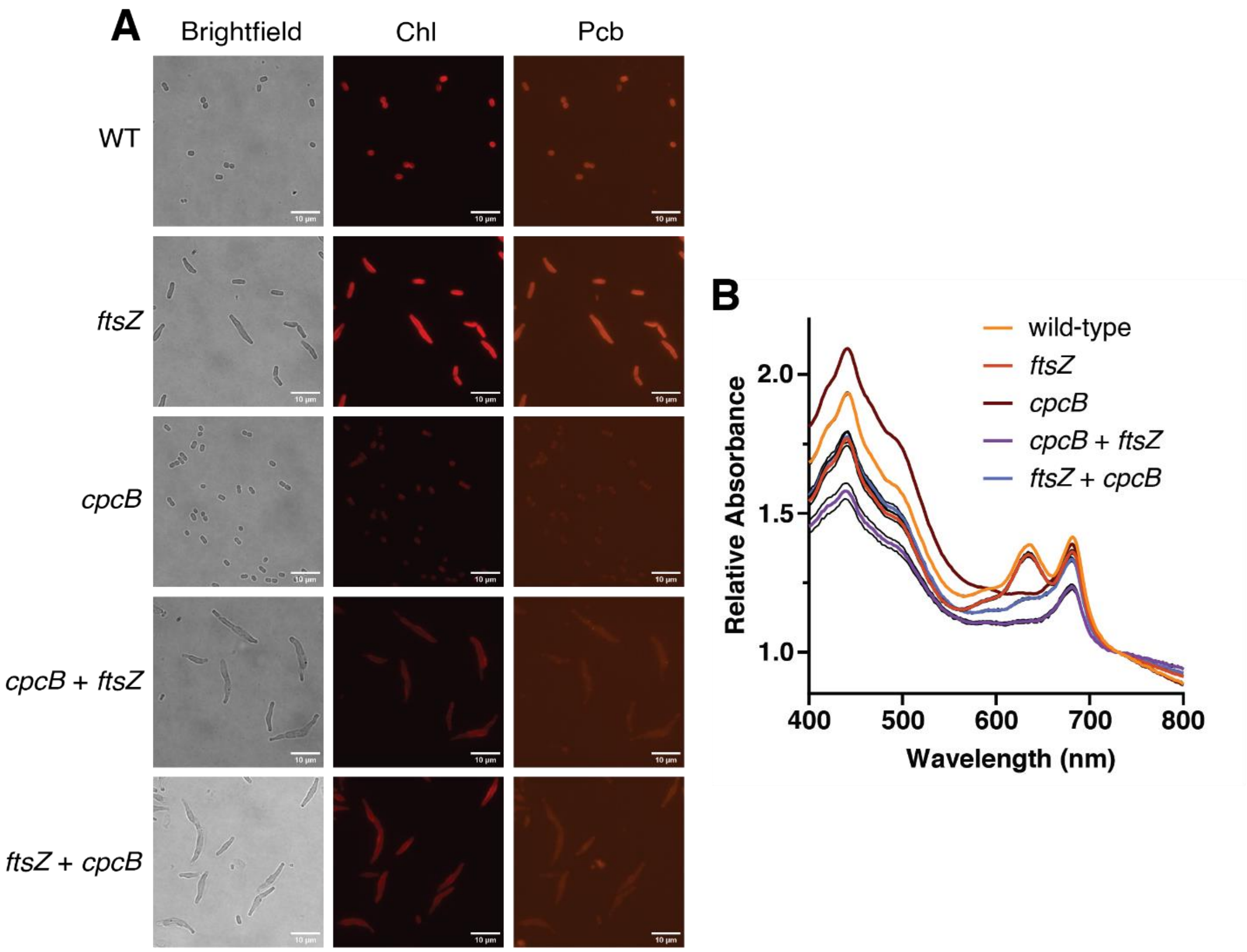


**Fig. S10. Dual guide constructs display combined knockdown phenotypes.** (A) We grew cells overnight in A+ media with inducers, expressing either no sgRNA (wild-type, WT), single sgRNAs, or double sgRNAs, then fixed them on a 1% agarose pad, imaged in sets of three. *ftsZ* repression led to elongated cells, while *cpcB* repression reduced phycobilin (Pcb) fluorescence. (B) We measured absorbance scans of each culture after overnight growth with induction using a Tecan Spark microplate reader. Indeed, they generated both phenotypes, regardless of sgRNA order. *cpcB* repression resulted in the characteristic reduction in Pcb absorbance at 630 nm. Error bars display the standard deviation from n=3 technical replicates.

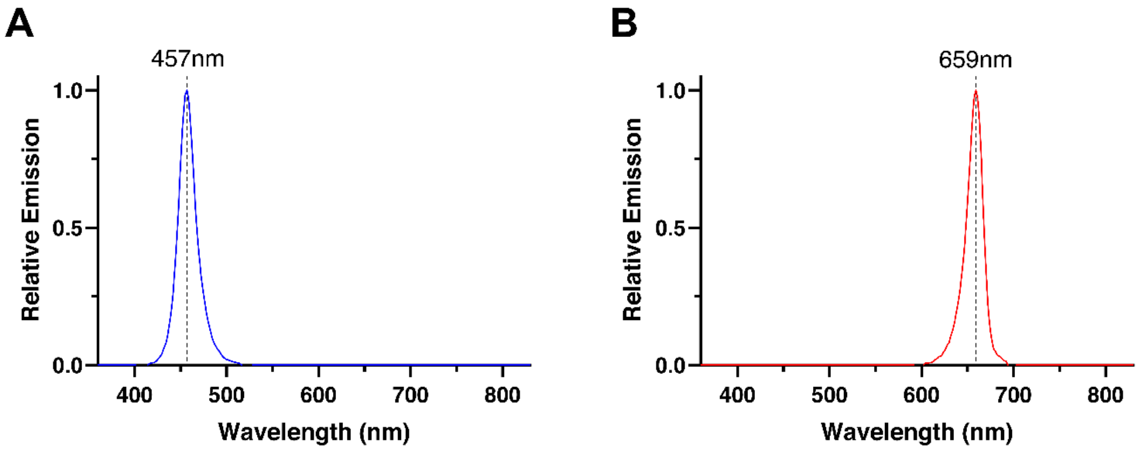


**Fig. S11. LEDs emit monochromatic blue or red light.** Absorption spectra of (A) blue and (B) red LEDs used in this experiment (HIGROW 36W LED light bulbs). We measured spectra using a Gigahertz-Optik MSC15 Spectral Light Meter.

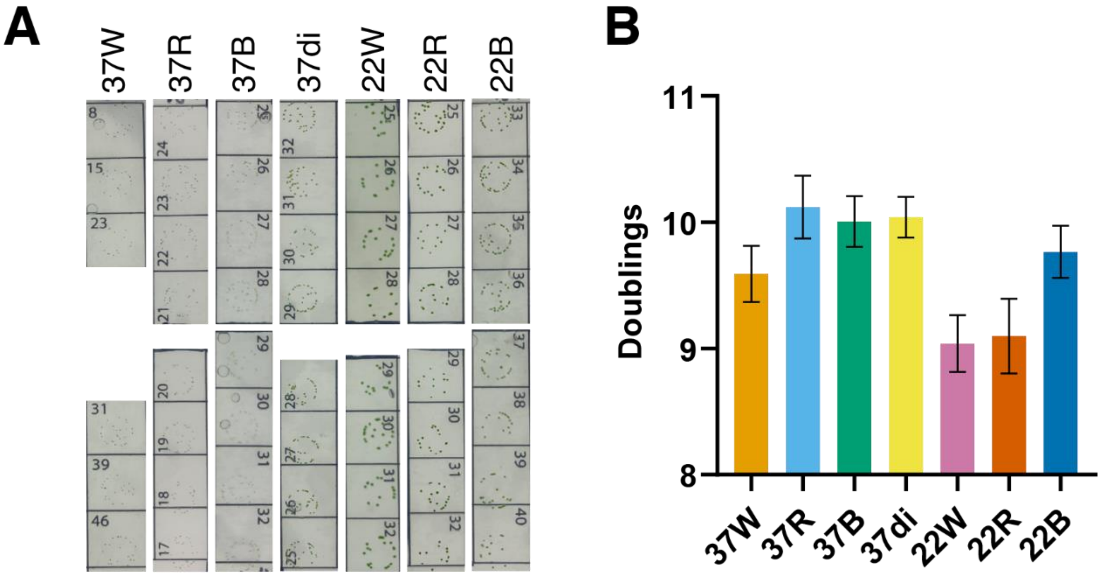


**Fig. S12. Libraries underwent a minimum of nine doublings within each library condition.** We retained a small volume of culture for doubling calculations from each genome-scale screen upon harvest (once both replicates exceeded an $OD_{730}$ of 0.3). (A) We diluted cultures by a factor of $10^4$ then spotted 5 µl with 3-4 technical replicates, counting colonies after incubation to calculate total CFUs. (B) We calculated doublings as the base 2 logarithm of the final CFU count divided by the initial CFU count in the inoculum. Data in (B) represent the mean and SEM of n=2 biological library replicates.

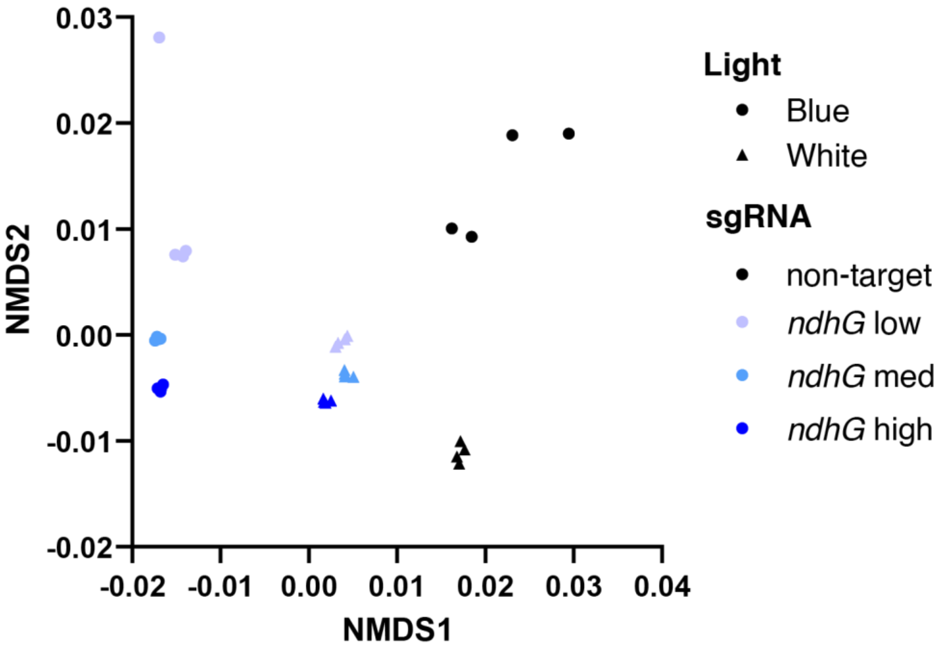


**Fig. S13. Proteomics replicates show strong agreement in expression levels.** We used non-metric multidimensional scaling (NMDS), a non-linear dimensionality reduction approach, to visualize similarity between proteomics samples. We observed tight clustering of replicates, indicating high similarity in protein expression levels. Data depict n=4 biological replicates per condition.

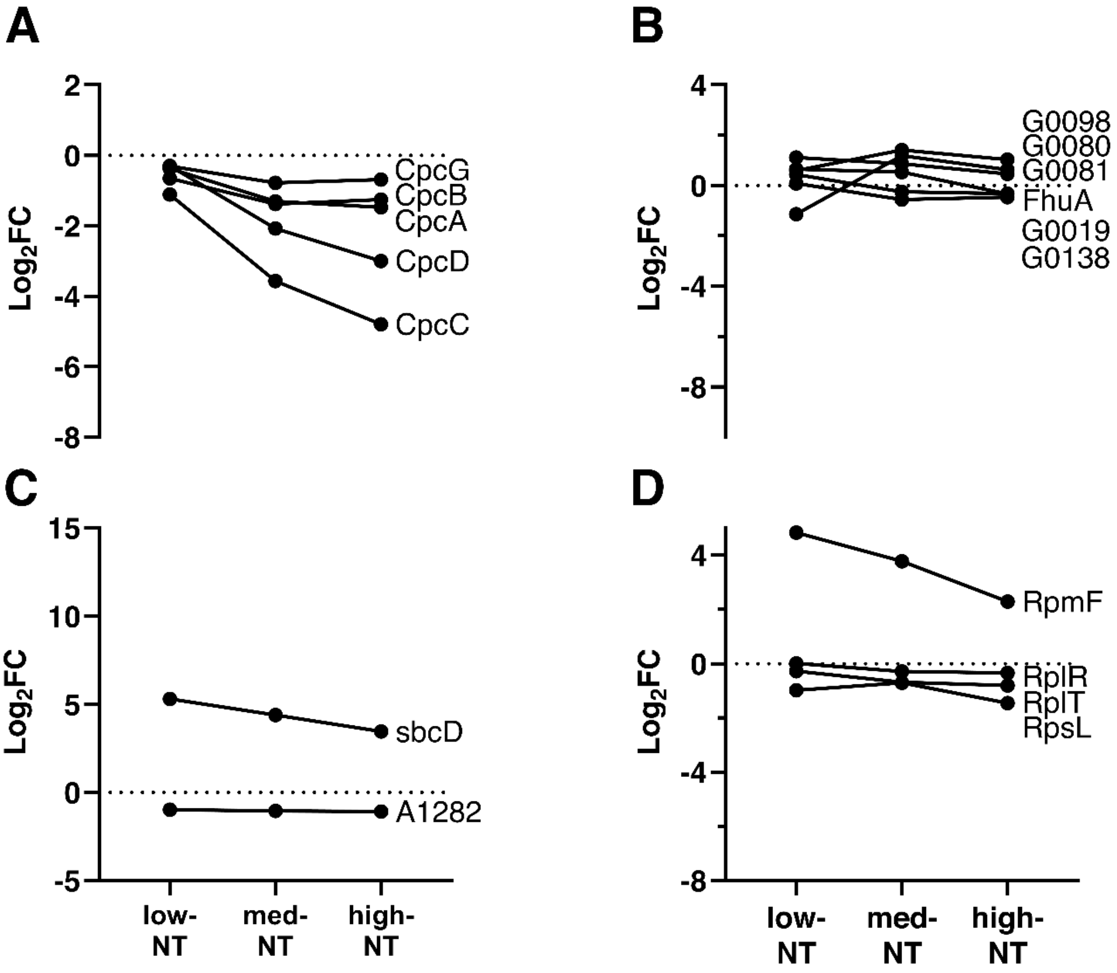


**Fig. S14. Protein levels respond differently to *ndhG* repression in 22W versus 22B.** (A) Cpc proteins exhibit a muted downward trend relative to 22B. (B) Iron handling and (C) proteins with increasing trends in 22B are not sensitive to *ndhG* knockdown. Proteins depicted in Figure 4 but not this figure were not detected in 22W. (D) Ribosomal proteins were either unaffected or enriched from *ndhG* knockdown, in contrast to the depletion observed in 22B.

**Table S1. Fold changes and raw read counts for sgRNAs across all growth conditions.** The accompanying Excel file provides the spacer sequences, DESeq2-calculated $Log_2FC$ values in each growth condition, the target gene, gene name, gene description, notes on protein homology with different cyanobacterial species, read counts, and DESeq2 statistical results for all sgRNAs in the CRISPRi library.

**Table S2. Proteomics results from samples grown in 22B and 22W.** The accompanying Excel file provides the raw protein abundance, $log_2$ abundance, ANOVA analysis of 22W samples and of 22B samples, and t-test results of each strain in 22W versus 22B.

**Table S3. Plasmids used in this study.**

| Plasmid name | Description | Source |
|---|---|---|
| pCas2F | Δ*acsA* TetR dCas9 plasmid (F RBS) -- dCas9 integration plasmid | Gordon (2016) Metabolic Engineering |
| KAMc0211 | ΔNS1_LacI_LacO-*minD* sgRNA_KmR -- IPTG inducible sgRNA expression | Moore (2019) BioRxiv |
| pAPH07 | glpK_LacI_LacO_BspQI-insertion_sgRNA_KmR -- single guide base integration plasmid | This work |
| pAPH021 | glpK_LacI_LacO_PaqCI-insertion_dual_sgRNA_KmR -- dual guide base integration plasmid | This work |
| p7-NT11 | glpK_LacI_LacO_nt-gRNA-11_sgRNA_KmR | This work |
| p7-ndhG-low | glpK_LacI_LacO_A0924_955778[c+]_sgRNA_KmR | This work |
| p7-ndhG-med | glpK_LacI_LacO_A0924_955949[c+]_sgRNA_KmR | This work |
| p7-ndhG-high | glpK_LacI_LacO_A0924_956008[c+]_sgRNA_KmR | This work |
| p7-gap | glpK_LacI_LacO_A0106_107358[c-]_sgRNA_KmR | This work |
| p7-psbB | glpK_LacI_LacO_A1759_1847732[c+]_sgRNA_KmR | This work |
| p7-psaC | glpK_LacI_LacO_A1589_1670525[c+]_sgRNA_KmR | This work |
| p7-trpS | glpK_LacI_LacO_A0037_35127[c-]_sgRNA_KmR | This work |
| p7-ftsZ | glpK_LacI_LacO_ A0024_21193[c-]_sgRNA_KmR | This work |
| p7-cpcB | glpK_LacI_LacO_ A2209_2300646[c-]_sgRNA_KmR | This work |
| p21-gap-psbB | glpK_LacI_LacO_A0106_107358[c-]_A1759_1847732[c+]_dual_sgRNA_KmR | This work |
| p21-psbB-gap | glpK_LacI_LacO_A1759_1847732[c+]_A0106_107358[c-]_dual_sgRNA_KmR | This work |
| p21-trpS-psaC | glpK_LacI_LacO_A0037_35127[c-]_A1589_1670525[c+]_dual_sgRNA_KmR | This work |
| p21-psaC-trpS | glpK_LacI_LacO_A1589_1670525[c+]_A0037_35127[c-]_dual_sgRNA_KmR | This work |
| P21-ftsZ-cpcB | glpK_LacI_LacO_A0024_21193[c-]_ A2209_2300646[c-]_dual_sgRNA_KmR | This work |
| P21-cpcB-ftsZ | glpK_LacI_LacO_ A2209_2300646[c-]_ A0024_21193[c-]_dual_sgRNA_KmR | This work |

**Table S4. Primers and oligos used in this study.**

| Oligo name | Sequence | Description |
|---|---|---|
| APH069 | acagctagaaatgaactcacctc | A2209_2300646[c-] Fwd, *cpcB* pAPH07 insert |
| APH070 | aacgaggtgagttcatttctagc | A2209_2300646[c-] Rev, *cpcB* pAPH07 insert |
| APH187 | acaATTCATATTATACCCTATCC | nt-gRNA_11 Fwd, non-target guide (NT11), p7-insert |
| APH188 | aacGGATAGGGTATAATATGAAT | nt-gRNA_11 Rev, non-target guide (NT11), p7-insert |
| APH287 | TCTGAGATGAGTTTTTGTTCGGGC | F primer, 7002 NGS diversification set |
| APH288 | NTCTGAGATGAGTTTTTGTTCGGGC | F primer, 7002 NGS diversification set |
| APH289 | NNTCTGAGATGAGTTTTTGTTCGGGC | F primer, 7002 NGS diversification set |
| APH290 | NNNTCTGAGATGAGTTTTTGTTCGGGC | F primer, 7002 NGS diversification set |
| APH291 | ACAGACATAAGTCCCATCACCGT | R primer, 7002 NGS diversification set |
| APH292 | NACAGACATAAGTCCCATCACCGT | R primer, 7002 NGS diversification set |
| APH293 | NNACAGACATAAGTCCCATCACCGT | R primer, 7002 NGS diversification set |
| APH294 | NNNACAGACATAAGTCCCATCACCGT | R primer, 7002 NGS diversification set |
| APH321 | acaCCCTGACGAAGGAGACGACC | A0924_956008[c+] Fwd, *ndhG* high repression (ndhG high), p7-insert |
| APH322 | aacGGTCGTCTCCTTCGTCAGGG | A0924_956008[c+] Rev, *ndhG* high repression (ndhG high), p7-insert |
| APH323 | acaTTACCAAAACCATCGTGCCA | A0924_955949[c+] Fwd, ndhG medium repression (ndhG med), p7-insert |
| APH324 | aacTGGCACGATGGTTTTGGTAA | A0924_955949[c+] Rev, ndhG medium repression (ndhG med), p7-insert |
| APH325 | acaTCTCAGGAATAATATCGCGA | A0924_955778[c+] Fwd, ndhG low repression (ndhG low), p7-insert |
| APH326 | aacTCGCGATATTATTCCTGAGA | A0924_955778[c+] Rev, ndhG low repression (ndhG low), p7-insert |

| Oligo name | Sequence | Description |
|---|---|---|
| APH355 | CTTAGAACTGCACCTGCTATA | Fwd – pAPH021 central insert amplification |
| APH356 | GGGTATAATATGAATCACCTGCATAT | Rev – pAPH021 central insert amplification |
| APH357 | cacaCATGATCGGAAATTGCGACG | A0024_21193[c-] p21-*ftsZ*-site1-fwd |
| APH358 | aaacCGTCGCAATTTCCGATCATG | A0024_21193[c-] p21-*ftsZ*-site1-rev |
| APH361 | gttcCATGATCGGAAATTGCGACGgtt | A0024_21193[c-] p21-*ftsZ*-site2-fwd |
| APH364 | ctaaaacCGTCGCAATTTCCGATCATG | A0024_21193[c-] p21-*ftsZ*-site2-rev |
| APH395 | cacaCGACGCCCCACTCTTTCCAA | A0106_107358[c-] p21-gap-low-site1-F |
| APH396 | aaacTTGGAAAGAGTGGGGCGTCG | A0106_107358[c-] p21-gap-low-site1-R |
| APH401 | gttcCCTGCCGCCACATGGGATTGgtt | A1759_1847732[c+] p21-psbB-low-site2-F |
| APH402 | ctaaaacCAATCCCATGTGGCGGCAGG | A1759_1847732[c+] p21-psbB-low-site2-R |
| APH407 | cacaGCACTTTATCGGCATCATAC | A0037_35127[c-] p21-trpS-low-site1-F |
| APH408 | aaacGTATGATGCCGATAAAGTGC | A0037_35127[c-] p21-trpS-low-site1-R |
| APH413 | gttcAACCATCTCTAGGACATCAAgtt | A1589_1670525[c+] p21-psaC-low-site2-F |
| APH414 | ctaaaacTTGATGTCCTAGAGATGGTT | A1589_1670525[c+] p21-psaC-low-site2-R |
| APH429 | acaCGACGCCCCACTCTTTCCAA | A0106_107358[c-] p7-gap-low-F |
| APH430 | aacTTGGAAAGAGTGGGGCGTCG | A0106_107358[c-] p7-gap-low-R |
| APH435 | acaGCACTTTATCGGCATCATAC | A0037_35127[c-] p7-trpS-low-F |
| APH436 | aacGTATGATGCCGATAAAGTGC | A0037_35127[c-] p7-trpS-low-R |
| APH441 | acaCCTGCCGCCACATGGGATTG | A1759_1847732[c+] p7-psbB-low-F |
| APH442 | aacCAATCCCATGTGGCGGCAGG | A1759_1847732[c+] p7-psbB-low-R |
| APH447 | acaAACCATCTCTAGGACATCAA | A1589_1670525[c+] p7-psaC-low-F |
| APH448 | aacTTGATGTCCTAGAGATGGTT | A1589_1670525[c+] p7-psaC-low-R |
| APH480 | cacaAACCATCTCTAGGACATCAA | A1589_1670525[c+] p21-psaC-low-site1-F (p21-blue-low-rev) |
| APH481 | aaacTTGATGTCCTAGAGATGGTT | A1589_1670525[c+] p21-psaC-low-site1-R (p21-blue-low-rev) |

| Oligo name | Sequence | Description |
|---|---|---|
| APH482 | gttcGCACTTTATCGGCATCATACgtt | A0037_35127[c-] p21-trpS-low-site2-F (p21-blue-low-rev) |
| APH483 | ctaaaacGTATGATGCCGATAAAGTGC | A0037_35127[c-] p21-trpS-low-site2-R (p21-blue-low-rev) |
| APH484 | cacaCCTGCCGCCACATGGGATTG | A1759_1847732[c+] p21-psbB-low-site1-F (p21-red-low-rev) |
| APH485 | aaacCAATCCCATGTGGCGGCAGG | A1759_1847732[c+] p21-psbB-low-site1-R (p21-red-low-rev) |
| APH486 | gttcCGACGCCCCACTCTTTCCAAgtt | A0106_107358[c-] p21-gap-low-site2-F (p21-red-low-rev) |
| APH487 | ctaaaacTTGGAAAGAGTGGGGCGTCG | A0106_107358[c-] p21-gap-low-site2-R (p21-red-low-rev) |
| APH547 | acaCATGATCGGAAATTGCGACG | A0024_21193[c-] fwd *ftsZ* p7 insert |
| APH548 | aacCGTCGCAATTTCCGATCATG | A0024_21193[c-] rev *ftsZ* p7 insert |
| APH549 | cacaGCTAGAAATGAACTCACCTC | A2209_2300646[c-] p21-*cpcB*-site1-fwd |
| APH550 | aaacGAGGTGAGTTCATTTCTAGC | A2209_2300646[c-] p21-*cpcB*-site1-rev |
| APH551 | gttcGCTAGAAATGAACTCACCTCgtt | A2209_2300646[c-] p21-*cpcB*-site2-fwd |
| APH552 | ctaaaacGAGGTGAGTTCATTTCTAGC | A2209_2300646[c-] p21-*cpcB*-site2-rev |

**Table S5. Cluster assignments of genes.** The accompanying Excel file contains the mean locus sgRNA $Log_2FC$ and assigned cluster category for all genes which contained at least one sgRNA with statistically significant enrichment or depletion.

**Table S6. Genes mentioned in this study.**

| Gene name | Locus | Gene description |
|---|---|---|
| *apcB* | A1929 | Allophycocyanin, beta subunit |
| *apcE* | A2009 | Phycobilisome core-membrane linker phycobiliprotein |
| *atpD* | A0749 | ATP synthase beta chain |
| *chlH* | A1000 | Magnesium-chelatase, subunit H |
| *chlP* | A2476 | Geranylgeranyl reductase |
| *cpcA* | A2210 | Phycocyanin, alpha subunit |
| *cpcB* | A2209 | Phycocyanin, beta subunit |

| Gene name | Locus | Gene description |
|---|---|---|
| *cpcC* | A2211 | Phycocyanin-associated rod linker protein |
| *cpcD* | A2212 | Phycocyanin-associated, rod-terminating linker protein |
| *cpcG* | A0811 | Phycobilisome rod-core linker polypeptide |
| *cpcT* | A2095 | Phycocyanobilin lyase |
| *crtR* | A0915 | Beta-carotene oxygenase |
| *cysS* | A2873 | Cysteinyl-tRNA synthetase |
| *desA* | A2756 | Phosphatidylcholine desaturase |
| *desC* | A2198 | Delta-9 acyl-lipid desaturase |
| *efp* | A0053 | Translation elongation factor P |
| *fhuA* | G0103 | Ferrichrome-iron receptor; TonB-dependent siderophore receptor |
| *fruB* | A2638 | Putative coenzyme F420 hydrogenase/dehydrogenase beta subunit |
| *ftsZ* | A0024 | Cell division protein FtsZ |
| *gap* | A0106 | -, type I |
| *glgC* | A0095 | Glucose-1-phosphate adenylyltransferase |
| *hliA* | A0858 | High light inducible protein hli5 |
| *n/a* | A0186 | CAB/ELIP/HLIP superfamily |
| *kaiA* | A0289 | Circadian clock protein |
| *kaiC* | A0287 | Circadian clock protein |
| *n/a* | A1282 | Conserved hypothetical protein |
| *n/a* | G0019 | Siderophore biosynthesis protein, lucA/lucC family |
| *n/a* | G0080 | Iron compound ABC transporter, ATP-binding protein |
| *n/a* | G0081 | Outer membrane TonB-dependent hemoglobin/transferrin/lactoferrin receptor family protein |
| *n/a* | G0098 | TonB-dependent siderophore receptor |
| *n/a* | G0138 | TonB-dependent siderophore receptor |
| *n/a* | F0018 | Uncharacterized protein |
| *ndhA* | A0926 | NADH dehydrogenase subunit A |
| *ndhD4* | A1806 | NADH dehydrogenase subunit D4 |
| *ndhF4* | A1805 | NADH dehydrogenase subunit F4 |
| *ndhG* | A0924 | NADH dehydrogenase subunit G |
| *petD* | A0841 | Cytb6/f complex subunit IV |
| *pgk* | A1585 | Phosphoglycerate kinase |
| *pmgA* | A0556 | Photomixotrophic growth related protein |
| *ppnK* | A0217 | ATP-NAD kinase, putative |
| *prk* | A2665 | phosphoribulokinase |
| *prs* | A2726 | Ribose-phosphate pyrophosphokinase |
| *psaC* | A1589 | Photosystem I iron-sulfur center subunit VII |
| *psaD* | A0682 | Photosystem I subunit II |
| *psbB* | A1759 | Photosystem II CP47 reaction center protein |
| *psbD* | A1560 | Photosystem II D2 protein |
| *psbF* | A0231 | Cytochrome b559, beta subunit |
| *psbH* | A0808 | Phosphoprotein of photosystem II |

| Gene name | Locus | Gene description |
|---|---|---|
| *purD* | A1406 | Phosphoribosylamine-glycine ligase |
| *purF* | A0550 | Amidophosphoribosyltransferase |
| *rbpA* | A1629 | RNA-binding protein |
| *rpaA* | A1308 | Two-component response regulator |
| *rpiA* | A1269 | Ribose 5-phosphate isomerase A |
| *rplR* | A1050 | Ribosomal protein L18 |
| *rplT* | A2180 | Ribosomal protein L20 |
| *rpmF* | A1893 | Ribosomal protein L32 |
| *rpsL* | A2064 | 30S ribosomal protein S12 |
| *rpoA* | A1042 | DNA-directed RNA polymerase, alpha subunit |
| *sasA* | A2818 | Adaptive-response sensory-kinase sasA |
| *sbcD* | A2342 | Probable exonuclease |
| *trpS* | A0037 | Tryptophanyl-tRNA synthetase |

**Table S7. sgRNA spacers used in this study.** Targeting guide IDs specify locus, base position of target within the genome, coding (c) or noncoding (nc) strand of locus, and plus (+) or minus (-) strand.

| ID | Description | Spacer sequence (5' – 3') |
|---|---|---|
| nt-gRNA_11 | Non-targeting sgRNA | ATTCATATTATACCCTATCC |
| A0924_955778[c+] | *ndhG* low repression | TCTCAGGAATAATATCGCGA |
| A0924_955949[c+] | *ndhG* medium repression | TTACCAAAACCATCGTGCCA |
| A0924_956008[c+] | *ndhG* high repression | CCCTGACGAAGGAGACGACC |
| A0106_107358[c-] | *gap* max enrichment | CGACGCCCCACTCTTTCCAA |
| A1759_1847732[c+] | *psbB* max enrichment | CCTGCCGCCACATGGGATTG |
| A1589_1670525[c+] | *psaC* max enrichment | AACCATCTCTAGGACATCAA |
| A0037_35127[c-] | *trpS* max enrichment | GCACTTTATCGGCATCATAC |
| A0024_21193[c-] | ftsZ repression phenotype | CATGATCGGAAATTGCGACG |
| A2209_2300646[c-] | *cpcB* repression phenotype | GCTAGAAATGAACTCACCTC |
